# HDI-STARR-seq: Condition-specific enhancer discovery in mouse liver in vivo

**DOI:** 10.1101/2024.06.10.598329

**Authors:** Ting-Ya Chang, David J Waxman

## Abstract

STARR-seq and other massively-parallel reporter assays are widely used to discover functional enhancers in transfected cell models, which can be confounded by plasmid vector-induced type-I interferon immune responses and lack the multicellular environment and endogenous chromatin state of complex mammalian tissues. Here, we describe HDI-STARR-seq, which combines STARR-seq plasmid library delivery to the liver, by hydrodynamic tail vein injection (HDI), with reporter RNA transcriptional initiation driven by a minimal *Albumin* promoter, which we show is essential for mouse liver STARR-seq enhancer activity assayed 7 days after HDI. Importantly, little or no vector-induced innate type-I interferon responses were observed. Comparisons of HDI-STARR-seq activity between male and female mouse livers and in livers from males treated with an activating ligand of the transcription factor CAR (*Nr1i3*) identified many condition-dependent enhancers linked to condition-specific gene expression. Further, thousands of active liver enhancers were identified using a high complexity STARR-seq library comprised of ∼50,000 genomic regions released by DNase-I digestion of mouse liver nuclei. When compared to stringently inactive library sequences, the active enhancer sequences identified were highly enriched for liver open chromatin regions with activating histone marks (H3K27ac, H3K4me1, H3K4me3), were significantly closer to gene transcriptional start sites, and were significantly depleted of repressive (H3K27me3, H3K9me3) and transcribed region histone marks (H3K36me3). HDI-STARR-seq offers substantial improvements over current methodologies for large scale, functional profiling of enhancers, including condition-dependent enhancers, in liver tissue in vivo, and can be adapted to characterize enhancer activities in a variety of species and tissues by selecting suitable tissue- and species-specific promoter sequences.

## Introduction

Enhancers, classically defined as position-independent, positively acting *cis*-DNA regulatory elements, recruit DNA-binding proteins and increase transcription through long distance interactions that bring them into close proximity with promoters of target genes [1]. Enhancer activation and cooperativity are key features governing cell-type specific gene expression [1, 2]. Genomic regions containing enhancers are often found at nucleosome-depleted (open chromatin) regions flanked by chromatin enriched for various combinations of activating histone marks (e.g., H3K27ac, H3K4me1, H3K4me3) and depleted of repressive histone marks (e.g., H3K27me3, H3K9me3) [3–7]. The pattern of histone marks is an important feature of the underlying chromatin state and is thought to directly impact chromatin accessibility and hence the availability of DNA regulatory elements for transcription factor (TF) binding [8]. Changes in chromatin state can be induced by both endogenous and environmental factors and may be retained in either short-term or long-term epigenetic memory, including transgenerational effects via the germ line [9–11].

Epigenetic features defining the chromatin state can be discovered genome-wide using chromatin immunoprecipitation (ChIP-seq) and nuclease accessibility assays (e.g., DNase-seq, ATAC-seq), which have identified tens of thousands of putative enhancer sequences, many of which are condition-dependent or cell type-specific [12–14]. Evolutionary conservation [12] and feature-based analysis methods for enhancer discovery [15, 16] have been described, but do not identify the full repertoire of active enhancers within the cell [4, 12, 17, 18]. Moreover, evolutionary sequence constraints alone do not determine when and where enhancers are active in vivo [12]. Enhancers can regulate genes via DNA looping over long distances, which further increases the challenge of enhancer discovery, given the vast and largely uncharacterized noncoding regions of the genome [19]. Tens of thousands of binding sites have been identified for many TFs by ChIP-seq, however, only a small fraction of such binding sites are thought to occur at active, functional enhancers [20]. These factors limit the utility of computational predictive approaches for enhancer discovery and have stimulated efforts to develop massively parallel reporter assays (MPRAs) for large-scale experimental screening of enhancer activity [21, 22].

STARR-seq (self-transcribed active regulatory regions sequencing) is a widely used transfection-based MPRA for the intrinsic transcriptional activity of any DNA fragment [23–25]. Typically, a library of DNA sequences comprising a set of putative enhancers is cloned into a STARR-seq reporter plasmid downstream of a minimal promoter and then transfected into a cell line of interest. DNA sequences that are active, functional enhancers drive the expression of a reporter RNA that incorporates the enhancer sequence itself. Deep sequencing of the corresponding cDNA both identities the enhancer sequence and gives a quantitative readout of its functional activity. STARR-seq is highly scalable and has been used to characterize enhancers that are functional in diverse biological contexts [26–28]. There are, however, several significant limitations to STARR-seq and other such MPRAs. First, STARR-seq has primarily been used to assay enhancer sequences in cultured cells, where the transfected enhancer DNA lacks its native chromatinized structure. A further concern is the absence of a native tissue microenvironment, with its extracellular matrix, secretory factors and myriad intercellular, cross-communication between the diverse cell types comprising complex mammalian tissues, which together have a profound impact on enhancer activity and gene expression [29, 30]. Moreover, serious complications, including large numbers of false positives and false negatives, can arise when the plasmid DNA vectors used for STARR-seq and other plasmid-based MPRAs activate strong innate immune responses in the transfected cells [25]. In vivo MPRA assays that may address some of these issues have been described, most notably for mouse brain using adeno-associated viral vectors [31–33], however, these viral vectors may introduce their own set of problems related to immunogenicity and toxicity [34, 35] leading to alterations in transcriptional regulatory networks and enhancer activity.

Tens of thousands of predicted enhancers have been identified in mouse liver as DNase-I hypersensitive sites (DHS) flanked by activating chromatin marks [36–38]. Many of these enhancer sequences show condition-specific chromatin accessibility, implicating them in key liver transcriptional processes, including responsiveness to drugs and environmental chemical exposures [36, 39] and sex-differences in gene expression [37, 40–42], which determine sex differences in liver disease susceptibility [43–45]. However, the functional analysis of these predicted enhancers has been hampered by the lack of a suitable large scale MPRA for mouse liver. Global analysis of liver enhancers has largely been limited to human HepG2 hepatoma cells [46–48], a transformed cell line that lacks the hepatocyte−non-parenchymal cell-cell interactions essential for normal liver function [49] and whose transcriptional and regulatory network features are a poor model for human liver [50]. Here, we address these issues, as well as the more general need for improved, non-viral methods to implement MPRA in live animals, by developing HDI-STARR-seq, which employs hydrodynamic injection (HDI) to deliver STARR-seq plasmid libraries to hepatocytes in an intact mouse liver model. We introduce a minimal *Albumin* promoter to the STARR-seq reporter library to enable expression in mouse liver, and we employ HDI to effect efficient, liver-specific delivery of plasmid DNA [51, 52] under conditions where hepatocytes, their cellular milieu and their interactions in three dimensions are preserved [53, 54]. We establish the liver promoter dependence of reporter activity, validate select results using conventional luciferase reporters delivered in vivo, and demonstrate the scalability of HDI-STARR-seq using both a small, focused reporter library and on a global scale, with a reporter library comprised of ∼50,000 genomic DNA fragments enriched for open chromatin regions. Finally, we demonstrate that liver STARR-seq enhancer activity is highly enriched at genomic regions in an active chromatin state in mouse liver and can be used to identify condition-specific regulatory elements linked to sex differences in liver gene expression and responsiveness to xenobiotic exposure. Thus, we show that HDI-STARR-seq is a highly effective method for global-scale characterization of regulatory sites in liver in vivo under a wide range of biological conditions.

## Methods

### Animal studies

Mouse work was carried out in accordance with ARRIVE Essential guidelines 2.0 [55] for study design, sample size, randomization, experimental animals and procedures and statistical methods, and with approval of the Boston University Institutional Animal Care and Use Committee (protocol # PROTO201800698). Young adult male and female CD-1 mice (ICR strain) were purchased from Charles River Laboratories (Wilmington, MA), housed in a temperature and humidity-controlled environment on a 12 h light cycle, and fed standard rodent chow. Reporter plasmids and STARR-seq plasmid libraries were delivered to the livers of male and female mice at 7-9 weeks of age by HDI on day 0, as detailed below. Mice were euthanized by cervical dislocation under CO_2_ anesthesia on day 7, unless stated otherwise. Livers were excised and flash frozen in liquid N2 and stored at -80°C. Livers were collected at a consistent time of day, between 10 AM and 12 noon (c.f., 7:30 AM lights on, 7:30 PM lights off), to minimize the impact of circadian rhythm on liver gene expression [56]. In some cases, a portion of the freshly excised liver was submerged in 4% paraformaldehyde for 48 h, then transferred to 70% ethanol for paraffin embedding and hematoxylin and eosin staining. Where indicated, HDI-injected mice were injected with TCPOBOP (Santa Cruz Biotechnology, Santa Cruz, CA, cat. #sc-203291), a CAR agonist ligand [57], at 3 mg TCPOBOP/kg body weight, or vehicle (control). TCPOBOP was dissolved in DMSO at 7.5 mg/ml, diluted 1:10 in corn oil, and then injected at 4 µl per g mouse body weight, with injections performed between 10:30-11:30 AM on day 6, i.e., 24 h prior to euthanasia on day 7.

### Plasmid delivery by HDI

Successful HDI requires tail vein injection of naked plasmid DNA in a large volume over a period of 5-8 sec [51, 52]. Plasmid DNA to be delivered to each mouse by HDI was resuspended in TransIT-EE delivery solution (MIR 5340, Mirus, Inc, Madison, WI) in a final volume of ∼3 ml, equivalent to 10% of the mouse body weight (vol/weight). For studies with a luciferase reporter readout, 15 μg of firefly reporter plasmid DNA (plasmids pTYC1-pTYC4; Table S1E) was mixed with 3 μg of Renilla reporter plasmid DNA (plasmid pGYL18; Table S1E). For STARR-seq studies, 15 μg of STARR-seq library plasmid DNA was mixed with 3 μg of Renilla reporter plasmid DNA per mouse. TransIT-EE was then mixed with the plasmid and kept at room temperature for ∼ 1 h to avoid cold shock during the injection. The plasmid + solution was transferred to a 5-ml syringe outfitted with a 27G, ½ inch needle (Fisher Scientific, cat. # 50-209-2293), taking care to not introduce any air bubbles. Mice were placed in a restraining tube and the tail vein was dilated using a glove filled with warm water. Each mouse was rapidly injected with a volume corresponding to ∼10% of its body weight, up a maximum of 3 ml (for mice > 30 g weight). HDI was typically completed within 5-8 sec and without appreciable resistance. HDI was largely ineffective, as judged by Renilla luciferase activity assayed in liver extracts (see below), when resistance to injection precluded delivery of the full volume or when the total time of injection exceeded 10 seconds.

### HDI transfection efficiency

The efficiency of plasmid delivery by HDI was determined by assaying Renilla luciferase assay, derived from the Renilla reporter plasmid pGYL18 included in each HDI transfection. A frozen piece of each liver (∼ 50 mg) was homogenized in 500 μl ice cold 1x Passive lysis buffer (Promega, cat. #E1910) using a rubber-tipped micro pestle fitting the bottom of a 1.7 ml centrifuge tube. A sample of the liver lysate (4 μl) was placed in each of 3 triplicate wells of a Greiner Bio 96 well white microplate (Greiner Bio, cat. #655207). Renilla luciferin reagent (25 μl of Stop & Glo Reagent, Promega Dual-Luciferase Reporter Assay System; Promega, Madison, WI, cat. #E1910) was injected into each well through one of the injectors of a CentroXS3 instrument (Berthold Technologies). Renilla luciferase activity was integrated over a 10 sec interval. Extracts were prepared from each set of livers in parallel to minimize batch effects. The absolute Renilla luciferase activity of each liver (mean value for triplicate assays from a single extract for each liver) was used to assess the success of HDI and to determine the mouse-to-mouse variation in liver HDI transfection efficiency. Firefly luciferase reporter plasmid activity (HDI experiments using plasmids pTYC1-pTYC4; Table S1E) was assayed in the same way but using 25 μl of LARII Reagent (Promega Dual-Luciferase Reporter Assay System, cat. #E1910) and 10 sec integrated intensity followed by Renilla luciferase assay determination in the same well. Reporter activity was normalized for HDI transfection efficiency by calculating the firefly luciferase activity/Renilla luciferase activity ratio for each liver (mean of technical triplicates for each activity assay) to obtain a quantitative readout of the activity of the cloned DNA sequence to be assayed. Successful HDI (>10-fold increase above a background luciferase activity of 100-135 activity units) was achieved in 23 of 36 mice (64%) injected with either a focused or a global STARR-seq plasmid library. Twenty of the 23 mice showed high liver Renilla luciferase activity (> 3,000 activity units) when assayed 7 days after HDI (Fig. S1B).

### Luciferase reporter plasmid construction

Genomic regions corresponding to individual DHS sites identified previously in vehicle control and TCPOBOP-treated adult male mouse liver [58] were amplified by PCR of CD-1 mouse liver genomic DNA and cloned upstream of the minimal *Alb* promoter sequence of KpnI/XhoI-digested plasmid pGYL1 [38] to generate plasmids pTYC1-pTYC4 (Table S1E). pGYL1 contains a 318 nt mouse *Alb* promoter sequence derived from pLIVE plasmid (Mirus Inc., Madison, WI, #MIR 5420) [59] followed by a 1,653 nt firefly luciferase reporter sequence derived from plasmid pGL4.10 (Promega). Similarly, the *Alb* enhancer-firefly luciferase reporter plasmid pGYL17 [38] was constructed by cloning into KpnI/XhoI-digested pGYL1 a 380 nt enhancer sequence located 10.5 kb upstream of the mouse *Alb* TSS, which corresponds to mouse chr5: 90,597,816-90,598,195 (C57BL/6J mouse genome, GRCm39). A corresponding Renilla luciferase reporter plasmid, designated pGYL18 (i.e., *Alb* enhancer-*Alb*-minimal promoter-Renilla luciferase), was also prepared (Table S1E).

### Construction of STARR-seq_Alb plasmid

Plasmids hSTARR-seq_SCP1 vector (Addgene plasmid # 99292, RRID:Addgene_99292; listed as pTYC6 in Table S1E) and hSTARR-seq_ORI vector (Addgene plasmid # 99296, RRID:Addgene_99296; pTYC7 in Table S1E) were gifts from Alexander Stark [25]. hSTARR-seq_SCP1 was modified to generate plasmid pTYC10 by removing the SCP1 promoter sequence by BanII digestion and replacing it with a minimal mouse *Alb* promoter sequence, comprised of 318 nt obtained from pLIVE plasmid (cat. #MIR 5420, Mirus Inc, Madison, WI), where the *Alb* promoter HNF1 TF binding site was changed from GATC to AATC to prevent bacterial methylation and to enable expression of the full intrinsic activity of the *Alb* promoter in HDI-transfected livers [59]. Oligonucleotides ON8196 and ON8197 (Table S1B) were used to attach homology arms and amplify the *Alb* promoter from the pLIVE-derived plasmid pGYL17 (see above). The 318 nt sequence of *Alb* promoter is as follows: TGCTCAAATGGGAGACAAAGAGATTAAGCTCTTATGTAAAATTTGCTGTTTTACATAACTTTAATGAATGGACAAAGTCTTGTGCATGGGGGTGGGGGTGGGGTTAGAGGGGAACAGCTCCAGATGGCAAACATACGCAAGGGATTTAGTCAAACAACTTTTTGGCAAAGATGGTATGATTTTGTAATGGGGTAGGAACCAATGAAATGCGAGGTAAGTATGGTTAATAATCTACAGTTATTGGTTAAAGAAGTATATTAGAGCGAGTCTTTCTGCACACAGATCACCTTCCTATCAACCCCACTAGCCTCTGGCAAA.

This *Alb* promoter sequence is required for high level, sustainable expression of RNA transcripts derived from STARR-seq reporters introduced into hepatocytes by HDI.

### Focused STARR-seq plasmid library

We selected 100 genomic regions (Table S1C) representing 100 open chromatin sites (DHS sites) in mouse liver, many of which are condition-specific, i.e., they display chromatin accessibility that is sex-dependent [38] or altered (either increased or decreased) following exposure to the xenobiotic TCPOBOP [58]. Each DHS sequence was PCR amplified from mouse genomic DNA using specific PCR primers (Table S1D) [36, 58]. Individual PCR reactions were set up as follows. AccuPrime Taq DNA polymerase (Roche, cat # 12346-086) was used to amplify each selected DHS sequence in a 10 μl reaction following the manufacturer’s instructions and using these cycling conditions: 94^◦^C for 3 min, followed by 30 cycles of 94^◦^C for 15 sec, 52^◦^C for 30 sec, and 68^◦^C for 1 min/kb, with a final 10 min elongation step at 68^◦^C. Genomic DNA extracted from male CD-1 mouse liver was used as PCR template (100 ng DNA per 10 μl reaction). Crude PCR products from individual reactions were combined to give one of two different pools based on the apparent PCR product fragment sizes (pool 1: fragments > 600 nt; pool 2: ˂ 600 bp and fragments ˃ 600 bp) without adjusting for the fragment DNA concentration of each reaction. Each pool was purified using a QIAquick spin column (QIAgen, #28115) to remove salts and unused primers. To optimize performance, pooled PCR fragments > 600 bp in length (300 ng per tube x 2 tubes, each diluted with 100 μl of Promega RQ1 DNase 10x buffer (cat. # M6101)) were lightly digested with 0.2 μl of DNase-I (RQ1 RNase-Free DNase-I, Cat. # M6101, Promega) for 15 sec at 37^◦^C. Reactions were terminated by adding 150 μl of 50 mM EDTA, followed by heating for 10 min at 65^◦^C to obtain a population of DNA fragments 100-500 bp in length. This mixture of digested DHS fragments was combined with an equal amount of pool 2 DHS fragments (600 ng), which are comprised of smaller undigested fragments (< 600 bp), and then purified using a QIAquick spin column (QIAgen, #28115).

The combined pool of purified DHS fragments (∼1 μg) was end repaired and ligated to NEBNext adaptor (included in NEB cat. # E7645S) following the manufacturer’s instructions. KAPA HiFi HotStart ReadyMix (2X) (Roche, cat. #KK2601) was then used to amplify the Illumina adaptor sequence-ligated DNA fragments to attach homology arms (HA1 and HA2 in Fig. 2B; 15 bp at each end) to the pool of fragments in 4 parallel PCR reactions (PCR primers ON8352_F_infucloning and ON8353_R_infucloning, Table S1B) using these cycling conditions: 98^◦^C for 45 sec, followed by 8 cycles at 98^◦^C for 15 sec, 65^◦^C for 30 sec, and 72^◦^C for 45 sec, with a final elongation step for 60 sec at 72^◦^C. Next, for each focused STARR-seq library, we used a 2:1 molar ratio of insert to SalI/AgeI-digested pTYC10 (125 ng of pPromALB_STARRseq plasmid; Table S1E) in a single 10 μl In-Fusion HD reaction (Clontech, cat. # 639650) via the homology arms. The reaction products were pooled and precipitated using sodium acetate/ethanol, resuspended in 10 μl of 10 mM Tris HCl, pH 8.5 buffer and stored at -80^◦^C prior to use to increase transformation efficiency [60]. A 2.5 μl aliquot of the precipitated material was transformed into 20 μl of electrocompetent MegaX DH10B bacterial cells (ThermoFisher Scientific, cat. # EC0113) using a Gene Pulser Xcell Total System (Bio-Rad) and Gene Pulser Electroporation Cuvettes (0.1 cm gap, Bio-Rad; cat No. 1652089) under these conditions: 2 kV, 25 µF, 200 ohms. LB bacterial culture medium (1 L) was used to amplify each focused STARR-seq plasmid library, which was purified using PureLink™ Endotoxin-Free Maxi Plasmid Purification Kit (ThermoFisher Scientific, cat. #A31231) to obtain 2 mg of the final focused STARR-seq plasmid library, STARR-TYC3/7.

### Global STARR-seq plasmid libraries

DNA fragments released by DNase-I digestion of purified mouse liver nuclei were purified and then pooled from digestions representing the following four biological conditions: untreated adult male CD-1 mice, 27-h TCPOBOP-treated male CD-1 mice, untreated adult male C57BL/6 mice, and 3-h TCPOBOP-treated male C57BL/6 mice (samples generated previously, with ∼15-25% of reads localized to DHS peak regions [58]). The combined pool of DNase-released genomic DNA fragments was purified by two-tailed selection of SPRI beads to obtain fragments in the desired size range, as follows. First, 0.6x volume of SPRI beads was used to remove high molecular weight DNA; next, an additional 1.2x the original DNA volume was added (i.e., 120 μl of SPRI beads added to 200 μl of pooled DHS material (0.6x), followed by an additional 240 μl SPRI beads). A total of 98 ng DNase-I fragments, ranging from 100 to 500 bp in size, was then used as starting material in a single adapter ligation reaction using NEBNext Ultra II DNA library prep kit for Illumina (E7645S) according to the manufacturer’s instructions, but with the Illumina adaptor diluted 10-fold due to the low amount of starting material. KAPA HiFi HotStart ReadyMix was then used to amplify the Illumina adaptor sequence-ligated DNA fragments to attach homology arms (Fig. 2B) to the pool of fragments in 21 parallel PCR reactions, using the same primers and reaction conditions described above for the focused STARR-seq library. Next, we used a 2:1 molar ratio of insert to SalI/AgeI-digested vector (125 ng) and set up 6 parallel In-Fusion HD reactions via homology arms (10 μl each reaction) for each global STARR-seq library. The reaction products from the 6 separate reactions were pooled and precipitated using sodium acetate/ethanol, resuspended in 10 μl of 10 mM Tris HCl, pH 8.5 buffer and stored at -80^◦^C. A 2.5 μl aliquot of the precipitated material was transformed into 20 μl of electrocompetent MegaX DH10B bacterial cells, which were then amplified by growth in 1 L of LB bacterial culture medium, followed by purification to obtain ∼2 mg of endotoxin-free global STARR-seq library plasmid DNA, as described above for the focused STARR-seq library. It is critical to achieve a high transformation efficiency to maintain library complexity and quality, most importantly for the global STARR-seq library.

### Preparation of Illumina sequencing libraries for STARR-seq plasmid, transfected DNA and transcribed RNA reporters

The following methods, adapted from the published STARR-seq protocol [60], were used in a nested PCR amplification protocol to characterize the transcriptional activity of each DHS sequence (putative regulatory region) by HDI-STARR-seq. Three types of sequencing libraries were prepared, as follows. **1)** <u>Plasmid library preparation</u>: the batch of purified plasmid DNA that was injected into each mouse by HDI was directly used as a PCR template and amplified using primers ON8381/ON8382 (22 cycles for 70 sec at 72^◦^C), followed by a nested PCR reaction (5 cycles for 70 sec at 72^◦^C) with individual sample-barcoded i5 and i7 Illumina TruSeq primers (New England Biolabs NEBNext Multiplex Oligos for Illumina, cat. # E7600S; Table S1B) **2)** <u>DNA library preparation</u>: 200 mg flash frozen liver tissue collected 7 d after in vivo transfection by HDI was homogenized in 750 μl of QIAgen Buffer P1 using a rubber-tipped micro pestle and passed through a 20G needle three times to homogenize large tissue pieces. The extracted plasmid DNA was bound to a QIAprep spin column (QIAgen miniprep kit, cat. # 27106), eluted in 10 mM Tris HCl, pH 8.5 buffer and then used as a template in a PCR reaction with primers ON8381 and ON8382 (Table S1B, Fig. 2B) (22 cycles for 70 sec at 72^◦^C), followed by a nested PCR reaction (5 cycles for 70 sec at 72^◦^C) with individual sample-barcoded i5 and i7 primers, as above; and **3)** <u>Transcribed RNA (reporter sequences) library preparation</u>: 120 μg of total liver RNA was isolated from 60-100 mg liver tissue from each HDI-STARR-seq-treated mouse by TRIzol guanidinium thiocyanate-phenol-chloroform extraction then purified by polyA selection using oligo(dT) beads (New England Biolabs, cat. #E7490L). The purified RNA was digested with Turbo DNase-I (Invitrogen, cat. # AM2238) at 37°C for 30 min, reverse transcribed using primer ON8731 (Table S1B) for STARR-seq RNA first strand synthesis and Superscript III First-strand Synthesis SuperMix (Invitrogen, cat. # 18080-400) and then treated with 10 mg RNase A per ml (Fermentas, cat. # EN0531) for 1 h at 37^◦^C. The cDNA obtained was purified using 1.8x SPRI beads then used as a template in a PCR reaction using primers ON8379-ON8380 (22 cycles for 70 sec at 72^◦^C), followed by the same i5/i7 nested PCR reaction described above (5 cycles for 70 sec at 72^◦^C) (Fig. 2B). Primers ON8381 and ON8382 were designed to specifically amplify the STARR-seq plasmid DNA libraries by targeting the unspliced intron incorporated into the original plasmid and present in both the plasmid and DNA libraries but spliced out of the reporter RNA sequences. In contrast, primers ON8379 and ON8380 straddle the RNA splice junction and were designed to specifically amplify the transcribed and spliced RNA reporter molecules (Fig. 2B). Libraries were subject to Illumina multiplex sequencing (Novogene, Sacramento, CA) (Table S3A).

### Analysis of Illumina sequencing data

Paired-end sequence reads, each 150 base pairs long, were generated for the STARR-seq plasmid, DNA and RNA libraries. Raw FASTQ files were input to a custom pipeline that produces quality control metrics, including FASTQC reports (FASTX Toolkit, version 0.0.13.2), identifies and removes contaminating adapter sequences using Trim_galore (version 0.4.2), and determines insert-size length distributions using Picard (version 1.123). Sequence reads were aligned to mouse genome mm9 using Bowtie2 (version 2.2.6) [61]. Peaks were identified in each STARR-seq library type (plasmid, DNA and RNA) for both the focused and the global STARR-seq libraries using MACS2 (Model-based Analysis of ChIP-Seq analysis, v2.1.0.20150731) with the option (–keepdup); all other parameters were set to default. The peak sets were filtered to eliminate peaks overlapping ENCODE blacklisted regions [62]. Peak sets for the focused libraries (Focus_MACS2_Peak_XXX; Table S1C) and separately for the global DHS libraries (G187_TCPO_RNA_XXX; Table S4A) were merged across all three types of libraries (plasmid, DNA and RNA libraries) and their biological replicates using the BEDTools merge command to generate a peak union list comprised of 1,673 MACS2 peaks (focused STARR-seq library) or 117,122 MACS2 peaks (global STARR-seq library). All genomic coordinates used in this study are based on mouse genome assembly mm9.

### Sequence read distributions and enhancer activity calculations for focused STARR-seq libraries

Sequencing depths are shown in Table S3A for each STARR-seq library (RNA, DNA and plasmid). 74-86% of the RNA reads in focused library sequencing samples G187M09-G187M16 mapped to the set of 1,673 MACS2 peaks; furthermore, the vast majority (89-93%) of RNA reads mapping to the 1,673 peak regions were localized to the 100 targeted genomic PCR regions (Table S3A). Furthermore, 65-70% of the DNA library and plasmid library sequence reads mapped to the 1,673 MACS2 peak regions, and 74% of those reads were within the 100 targeted genomic PCR regions (Table S3A). The other 1,573 peaks represent genomic regions amplified non specifically during the 100 targeted genomic-DNA PCR reactions and are sufficiently well represented to be detected as significant MACS2 peaks. The STARR-seq enhancer activity of each peak region was calculated as the ratio of RNA sequence reads mapping to that region to that of the extracted DNA library in the same condition (full dataset in Table S3B), as follows. All sequencing fragments that overlapped a MACS2 peak by at least 1 bp were included in the MACS2 peak region mapped reads reported in Table S3B. Raw sequence reads in each focused library (Table S3B) were normalized to the sequencing read depth based on the total number of reads in the 1,673 MACS2 peak regions (Table S3A). The mean enhancer activity of each MACS2 peak was equal to the normalized RNA reads in the peak region, averaged across all livers in a group, divided by the average DNA activity for that region. Fold change and p-values (Student’s t-Test, unpaired) for TCPOBOP-treated male livers (n=2), control male livers (n=2) and control female livers (n=4) were then calculated from the enhancer activities determined for the set of TCPOBOP-treated male livers as compared to the set of control male livers. These values represent the extent of increase or decrease in enhancer activity following TCPOBOP treatment. Similarly, sex differences in enhancer activity (fold change, p-values, Student’s t-Test, unpaired) were calculated for control female versus control male liver. Peak regions displaying condition-specific enhancer activity were identified at p-value < 0.05. Stringent condition-specific enhancers met the additional threshold of >2-fold difference in enhancer activity between biological conditions.

### Qualified genomic regions and enhancer activity determination for global STARR-seq libraries

Genomic regions represented in the global DHS STARR-seq libraries were qualified for inclusion in downstream analysis based on the number of Illumina sequence reads obtained from the original STARR-seq plasmid library, the number of sequence reads obtained from biological replicates for STARR-seq RNA extracted from each liver, and the consistency across biological replicates (Fig. S6B). First, we reduced the list of genomic regions under consideration from 117,122 merged MACS2 peaks (including 114,452 autosomal peaks) to 50,332 qualified autosomal genomic regions (mean width, 258 nt; Table S4A) by filtering out genomic regions with ≤ 40 sequencing depth-normalized Illumina sequence read pairs per 10 million mapped plasmid library STARR-TYC6 reads. Next, for each reporter RNA sequencing library prepared from RNA extracted from an HDI-transfected liver, BEDtools was used to count the number of mapped RNA sequence reads that overlapped the set of 50,332 qualified autosomal genomic regions. A qualified genomic region was designated an active enhancer if there were at least 20 normalized Illumina sequence reporter RNA library read counts per 10 million mapped reporter RNA reads across the region for one or more of the following: at least 3 of the 4 biological replicate livers from untreated male mice; all 3 biological replicates livers from untreated female mice; or at least 3 of the 4 biological replicates from TCPOBOP-treated male mice (Table S4C). A total of 8,857 active enhancers were identified, of which 1,556 passed the above thresholds for all 3 biological conditions tested (robust active enhancers). Genomic regions whose reporter RNA libraries gave < 20 normalized sequence read counts per 10 million mapped reporter RNA library reads in all 11 biological replicate livers derived from the above 3 biological conditions were designated stringent inactive enhancers (Table S4C). The mean enhancer activity of each of qualified genomic regions under each biological conditions was based on the mean normalized RNA library read counts for the corresponding set of livers, after excluding livers with < 20 normalized RNA sequence reads per 10 million mapped RNA reads to minimize artifactual results. Genomic regions that met the 20 normalized RNA sequence reads per 10 million mapped RNA read threshold for at least 3 out of 4 individual male livers, 3 out of 4 individual TCPOBOP-treated male livers, or 3 out of 3 individual female livers were designated active enhancer regions for that condition (Fig. S6B and Table S4C).

### Mapping STARR-seq reporter regions to gene targets

A single putative gene target (RefSeq or lncRNA gene) was assigned to each reporter sequence included in the STARR-seq libraries based on the closest gene transcription start site within the same TAD as the reporter sequence, as determined using BEDtools. Mapping results for the TCPOBOP-responsive genomic regions identified using the focused STARR-seq library are included in Table S3C, for sex-biased DHS in Table S3D and for the global STARR-seq library in Table S4C.

### Enrichment analysis

Various sets of active enhancers identified by HDI-STARR-seq were overlapped with published sets of DNase-seq peak regions (DHS) and ChIP-seq datasets for histone marks in male and female mouse liver [37, 40] and in control and TCPOBOP-exposed male mouse liver [36]. These analyses used a merged list of 83,422 mouse liver DHS (Table S4A), obtained by combining two published mouse liver DHS datasets [36, 37] using BEDtools. 14,330 of these DHS overlapped the set of 50,332 qualified genomic regions present in the global HDI-STARR-seq library. The HDI-STARR-seq enhancer sets were also overlapped with ChIP-seq peak sets for the growth hormone-regulated TFs STAT5 [63], Bcl6 [63] and Cux2 [64], and in the case of TCPOBOP-responsive enhancers, the nuclear receptor CAR [65]. Overlaps were also determined for chromatin state map assignments developed using a combination of DNase hypersensitivity and a panel of 6 histone marks determined separately for male and for female mouse liver [40]. When performing overlap analysis between HDI-STARR-seq active enhancers and specific histone marks, a 200 bp extension was added both upstream and downstream of the enhancer sequence to encompass histone modifications that flank enhancers centered at an open chromatin region. Detailed results are shown in Table S4E.

Enrichment scores (ES values) were calculated as follows. To determine the enrichment of a given HDI-STARR-seq enhancer set in mouse liver open chromatin (liver DHS peak regions), as compared to that of a background set comprised of the 7,787 HDI-STARR-seq stringently inactive genomic regions (Table S4C), we calculated ES = ratio A/ratio B, where ratio A = number of enhancer regions that overlap the liver DHS peak set, divided by the number of enhancer regions that do not overlap the liver DHS peak set; and ratio B = number of stringently inactive genomic regions that overlap the liver DHS peak set, divided by the number of stringently inactive genomic regions that do not overlap the liver DHS peak set. The significance of enrichment was determined by Fisher’s exact test, implemented in the R package stats v.3.6.0. For example, of 1,556 robust active enhancers identified by HDI-STARR-seq in all three biological conditions investigated (Fig. 5A), 1,341 enhancer regions overlap one or more DHS peaks identified in mouse liver and 215 regions do not overlap. Furthermore, of the 7,787 stringently inactive genomic regions, 901 regions overlap a DHS peak and 6,886 regions do not show overlap. Thus, ES = (1,341/901)/(215/6,886) = 47.7. The same approach to ES calculation was implemented for other sets of active enhancer regions, and for a set of 33,688 other qualified genomic regions from the global HDI-STARR-seq library that passed the threshold for enhancer activity in at least one of 11 of the individual livers tested but were not sufficiently reproducible between livers to meet the threshold for active enhancers.

ES values for various sets of HDI-STARR-seq enhancers for being enriched in a given histone mark dataset (e.g., enrichment of genomic regions with a particular histone mark in a given set of active enhancers, as compared to that of a background set comprised of 7,787 stringently inactive enhancers) were calculated as follows: ES = ratio A/ratio B, where ratio A = number of active enhancers that overlap the histone mark peak set, divided by the number of active enhancers that do not overlap the histone mark peak set; and ratio B = number of stringently inactive genomic regions that overlap the histone mark peak set, divided by the number of stringently inactive genomic regions that do not overlap the histone mark peak set. For example, of the 3,595 active enhancers identified in control female livers (Fig. 5A), 1,849 regions overlap an H3K27ac peak(s) identified in female liver and 1,746 regions do not overlap. Furthermore, of the 7,787 stringently inactive genomic regions, 478 regions overlap an H3K27ac peak found in control female liver, and 7,309 regions do not overlap. Thus, ES = (1,849/1,746)/(478/7,309) = 16.2.

### Association of active enhancers with genes showing TCPOBOP-responsiveness or sex-dependent expression

RefSeq and lncRNA genes showing significant changes in expression in male or female mouse liver 27-hr after TCPOBOP injection were identified from a list of 24,197 RefSeq genes and 3,152 multi-exonic lncRNA genes [66]. A total of 734 TCPOBOP-induced genes were identified and included in our analyses, based on these threshold values: fold change > 1.5 and adjusted p-value < 0.001 for RefSeq genes; and a fold change > 2 and adjusted p-value < 0.05 for lncRNA genes [66]. We also examined 983 liver-expressed genes (FPKM >1) that showed a significant sex-bias in expression (EdgeR adjusted p-value < 0.01), including 485 male-biased genes and 498 female-biased genes [67].

### Chromatin state analysis of STARR-seq regions in liver

Chromatin state maps previously defined for adult male liver, and separately, for adult female liver [40] using ChromHMM [8] were employed to determine the chromatin state enrichments for various sets of STARR-seq regions. These chromatin states are based on a 14-state model (states E1-E14) and were previously used to characterize sex differences in chromatin state and chromatin structure and their relationships to sex-biased gene expression [40]. Bedtools overlap was used to assign the chromatin state in each sex to each STARR-seq region. A STARR-seq region that overlapped a genomic region assigned to two or more different chromatin states was assigned the state with the longest overlap in bp. The enrichment, or depletion, of each chromatin state in the set of STARR-seq regions was graphed separately for male and female liver using bar plots.

### Positional enrichment of enhancer features at the center of active enhancers

We investigated the spatial distribution of enhancer features within active enhancer regions. To accomplish this, we analyzed three adjoining sub-regions, namely the left (5’ flank), middle, and right (3’ flank) subregions around the midpoint of each active enhancer, with each sub-region spanning a distance of 400 bp. We first calculated the midpoint of each enhancer and then expanded this region by 200 bp on each side, defining it as the middle subregion. The left subregion was then defined as the 400-bp region upstream of the middle subregion, and the right subregion defined as the 400-bp region downstream of the middle subregion. This approach allowed us to determine the relative positional enrichment of enhancer features within the context of active enhancers.

### Genome browser tracks

Illumina sequencing files and processed datasets are available at GEO under accession # GSE267205. Custom UCSC Genome Browser track files for the focused libraries investigating TCPOBOP responsiveness are available at: https://genome.ucsc.edu/s/Tingyac/G187_focus_RNA_group_cyp2b10_shiny. Custom UCSC Browser track files for the focused libraries investigating sex responsiveness are available at: https://genome.ucsc.edu/s/Tingyac/G187_focus_figS5. Normalized enhancer activity tracks (labeled ‘Norm. enh. activity’) were generated to quantify enhancer activity across various conditions. These tracks were computed using the WiggleTools package, utilizing bigwig files from both DNA and RNA libraries corresponding to the same conditions. DNA reads-normalized enhancer activity tracks included in these UCSC genome browser sessions were calculated as the ratio of normalized RNA sequence reads to normalized DNA sequence reads. Similarly, UCSC browser tracks showing enhancer responsiveness to TCPOBOP exposure and sex differences in enhancer activity were calculated as the ratio of DNA-normalized enhancer activity between biological conditions on a per DHS region basis.

## Results

### DHS reporter plasmid sequences delivered to mouse liver by HDI respond to xenobiotic treatment

We sought to develop an *in vivo* reporter assay for condition-specific enhancers identified as open chromatin regions (DHS) in mouse liver. We initially tested this approach by assaying a series of DHS sequences whose accessibility increases following treatment of mice with TCPOBOP, a selective agonist ligand of the nuclear receptor CAR (Nr1i3) [57], as determined by DNase-seq [36]. Five liver DHS sequences were cloned into the firefly luciferase reporter plasmid pPromALB_FireflyLuc (pGYL1; Table S1E), whose minimal *Alb* promoter sequence was modified to block bacterial methylation of a critical HNF1 binding site [38, 59]. We selected the minimal *Alb* promotor based on the high and liver-specific expression of *Alb* in mouse liver and its established ability to facilitate transgene expression in hepatocytes when transfected by HDI [59]. An *Alb* enhancer sequence 10.5 kb upstream of the mouse *Alb* gene [38, 68] served as a positive control for enhancer activity, and pPromALB without a DHS insert (empty vector pGYL1; Table S1E) defined the assay system’s baseline (background) activity. Plasmids were delivered to mouse liver by HDI, an established method for long-term, liver-specific transfection [51, 52] that can be used to assay DHS enhancer activity in mouse liver [38]. Mice were treated with a receptor-saturating dose of TCPOBOP given 6 days after HDI, i.e., after decay of the initial, high activity shown by the *Alb* minimal promoter [38, 59]. Livers were collected 24 h later (Fig. 1A), at which time exposure to TCPOBOP induces or represses >1000 genes in mouse liver [66]. Firefly luciferase activity was normalized to the activity of a co-transfected pPromALB-based Renilla luciferase reporter plasmid driven by the -10.5 kb *Alb* enhancer (pGYL18; Table S1E) to control for the substantial individual mouse-to-mouse variation in the efficiency of plasmid DNA delivery by HDI.

**Fig. 1.**
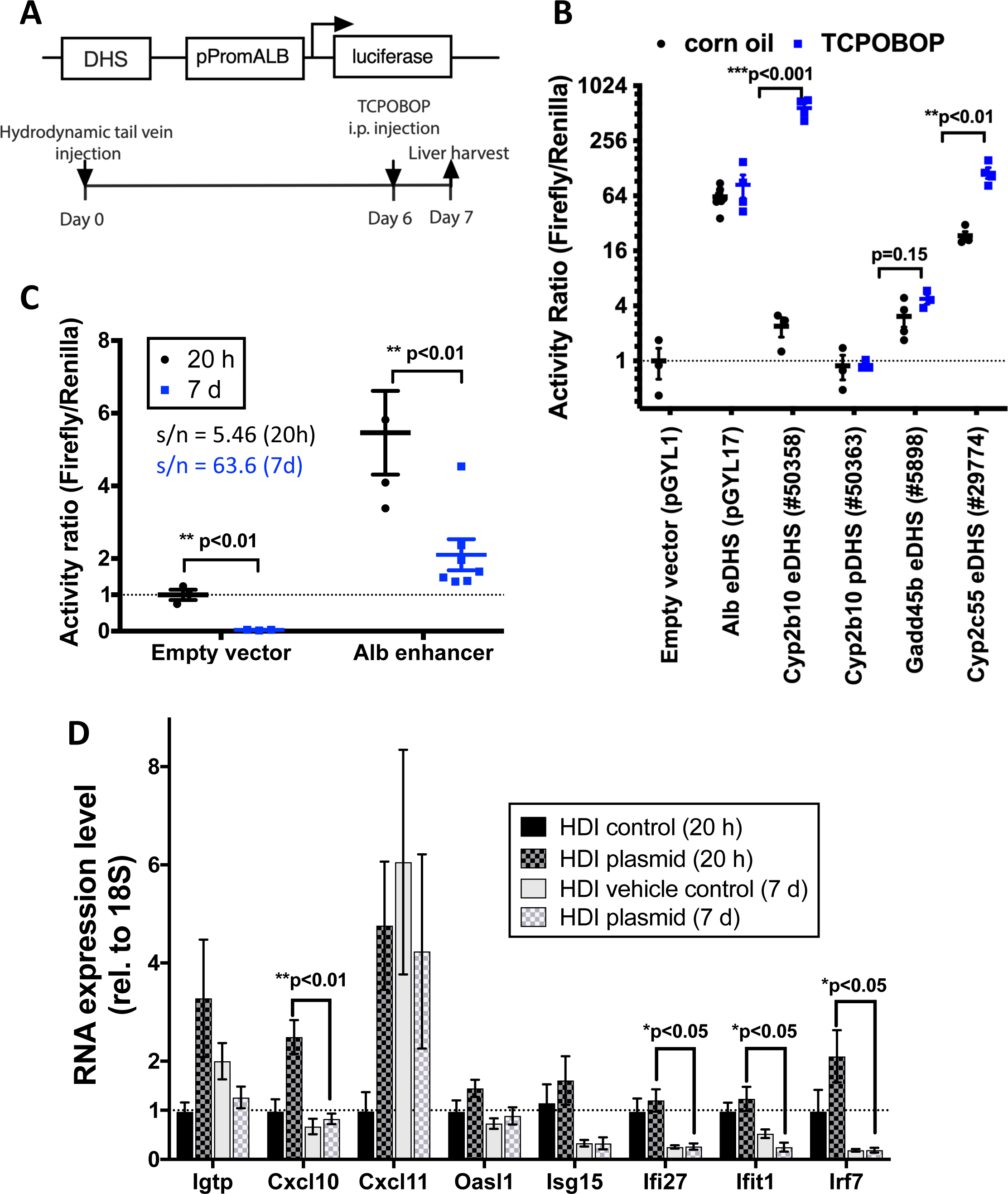
HDI-luciferase assay for liver DHS mapping to TCPOBOP-induced genes. (A) Mouse study design. Reporter plasmid encoding firefly luciferase driven by *Alb* minimal promoter, or by 1 of the 4 other individual DHS sequences indicated along the X-axis of panel B, was mixed with a normalization control plasmid encoding Renilla luciferase, then delivered to mouse liver by HDI on day 0, followed by TCPOBOP injection on day 6. (B) Normalized luciferase reporter activity for the DHS sequences shown along the X-axis, with TCPOBOP or corn oil (vehicle control) treatment (mean +/- SEM for n=3-7 individual livers per group; each symbol represents a single mouse liver). TCPOBOP-induced reporter activity was assessed by t-test: **, p <0.01; ***, p<0.001 for mean of corn oil control vs TCPOBOP group activity. The mean luciferase activity of the empty vector control group was set = 1 for normalization. “+”, TCPOBOP stimulates chromatin opening at the DHS [58]. (C) Impact of time after HDI (20 h vs. 7 d) on signal-to-noise ratio (S/N) for liver luciferase reporter activity. S/N was calculated as the ratio of normalized activity for *Alb* enhancer plasmid (pGYL17) vs empty vector plasmid (pGYL17) at each time point. Data shown are mean +/- SEM values, with the empty vector at 20 h group mean set = 1. *t*-test: **, p <0.01; n=3-7 per group. (D) qPCR analysis of interferon-related response genes showing the mean (+/- SEM) expression level of each gene in livers from mice treated by HDI to deliver plasmid DNA or vehicle control. Plasmids delivered by HDI (see Table S1E): HDI control = pGYL18 only; HDI plasmid = pGYL18 + pGYL17 or pGYL18 + pGYL1. Significance: *t*-test comparing livers with HDI delivery of reporter plasmid DNA and collected 20 h vs. 7 d later: *, p <0.05; **, p <0.01 for n=3-7 livers/group. See Table S1A for qPCR primers.

**Fig. 2.**
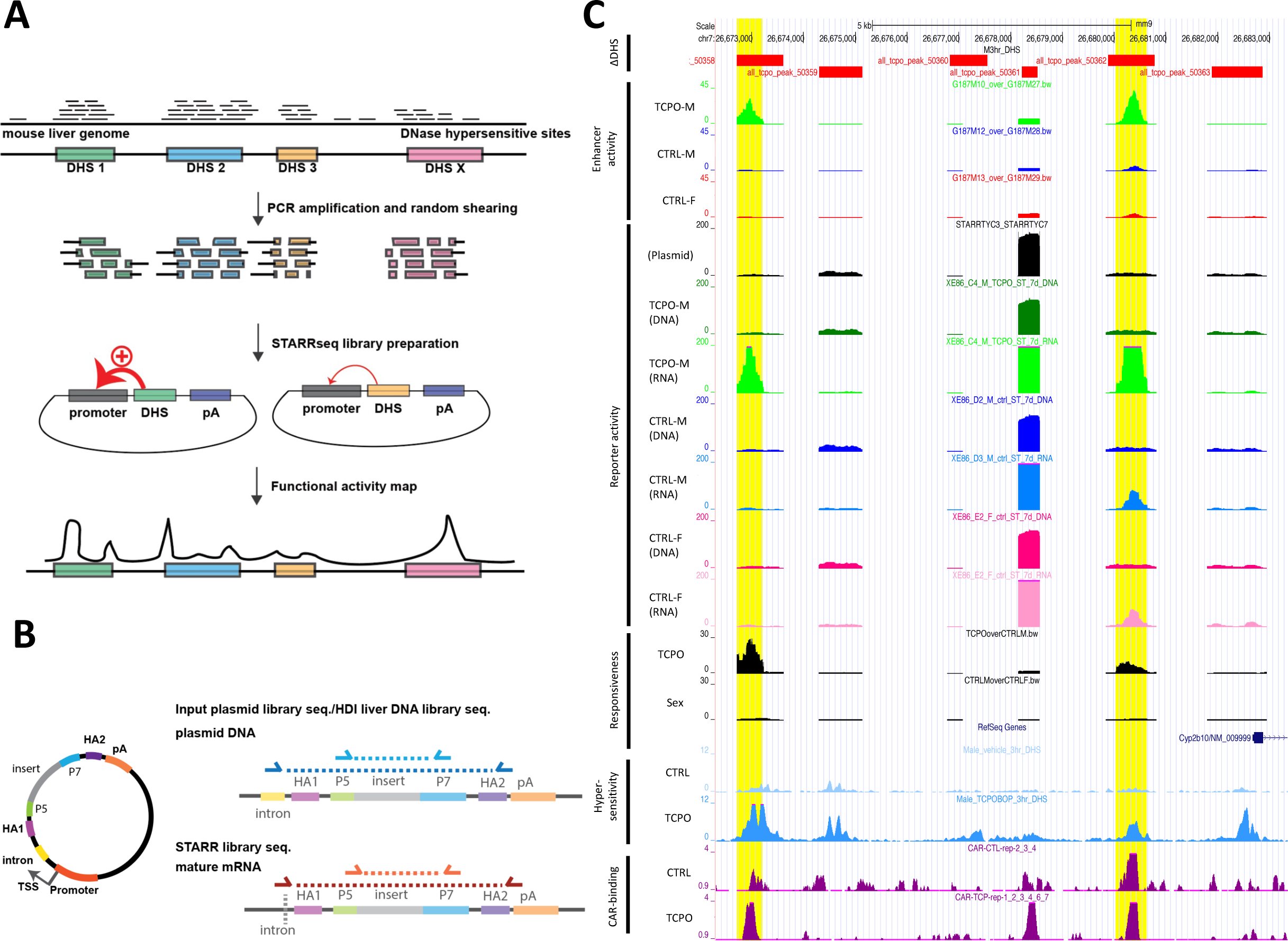
HDI-STARR-seq experimental design. (A) STARR-seq reporter plasmid design and library preparation. A pool of PCR-amplified and lightly digested DHS fragments released from mouse liver nuclei was cloned into the STARR-seq plasmid library pPromALB_STARR-seq (pTYC10), where they were inserted downstream of an *Alb* minimal promoter. Active enhancer DHS stimulate large increases in transcription of their own sequences, as determined by RNA-seq analysis of RNA extracted from HDI-transfected liver. Strong enhancers (+) can thus be distinguished from weak or inactive enhancers. Data analysis gives the functional activity map (bottom)), where the abundance of each transcribed sequence, detected as a peak of sequence reads mapped back to the mouse genome, indicates the intrinsic transcriptional activity of the DHS. (B) Plasmid map (circular STARR-seq reporter plasmid map) and linear maps of the resultant DNA and transcribed RNA reporters extracted from HDI-treated livers, both of which are present in the HDI-treated livers, but can be distinguished by the presence of a human intron sequence downstream of the promoter (top map, region shown in yellow) by using the indicated sets of PCR primers (Table S1B): mature RNA reporter molecules without the intron cannot be targeted by DNA reporter-specific primers after splicing. (C) UCSC Genome Browser view of focused STARR-seq library results for *Cyp2b10* region. Tracks show locations of the following elements (top to bottom): DHS regions where chromatin opens following TCPOBOP treatment (ΔDHS, red bars); normalized STARR-seq enhancer activity, calculated as the ratio of normalized RNA reads to normalized DNA reads for the 3 indicated conditions; and STARR-seq read pileups for the input plasmid library (Plasmid), the extracted DNA libraries (DNA) and the transcribed RNA libraries (RNA) from TCPOBOP-treated male livers (green), vehicle-treated male livers (blue) and vehicle-treated female livers (pink/red). Pairs of adjacent DNA and RNA tracks were for samples extracted the same HDI-treated mouse. Of the 6 DHS sites at or upstream of the *Cyp2b10* promoter, 5 DHS were well represented in the input plasmid library and HDI DNA library, and 2 DHS (DHS #50358, DHS #50362; yellow highlight) showed a strong increase in normalized enhancer activity with TCPOBOP treatment (second vs. third track from top). Enhancer responsiveness to TCPOBOP exposure, or regarding sex-specificity, is displayed in the last two tracks, and was calculated from the ratio of normalized enhancer activities for the corresponding pairs of conditions (TCPOBOP-exposed male vs. vehicle control male, and control male vs. control female).

TCPOBOP treatment (1 day exposure) significantly increased normalized firefly reporter activity for 2 of the 5 DHS sequences tested (Fig. 1B, Table S2A). DHS #50358, which is 9.5 kb upstream of the highly TCPOBOP-inducible gene *Cyp2b10* [66], showed a striking 245-fold increase in reporter activity following TCPOBOP treatment (normalized activity increased from 2.4 to 585, with empty vector control plasmid activity (plasmid pGYL1) set = 1.0). DHS #29774, located 1.5 kb upstream of the highly TCPOBOP-inducible gene *Cyp2c55*, showed a 9.8-fold higher basal reporter activity than DHS #50358, but a much smaller TCPOBOP-stimulated increase in activity (4.9-fold; normalized activity increased from 23 to 114). A high basal reporter activity but no increase after TCPOBOP treatment was observed for the -10.5 kb *Alb* enhancer (plasmid pGYL17); this is consistent with the unresponsiveness of *Alb* to TCPOBOP in mouse liver [66]. DHS #50363, which overlaps the *Cyp2b10* promoter, was inactive in both control and TCPOBOP-treated liver, indicating this genomic sequence is devoid of intrinsic enhancer activity. Finally, DHS #5898, a putative enhancer located 28 kb downstream of the TCPOBOP-inducible gene *Gadd45b*, and whose chromatin opens following TCPOBOP treatment [58], showed low basal activity (mean activity = 3.1 relative to empty vector control), and no significant increase following TCPOBOP treatment (mean activity = 4.8) (Fig. 1B, Table S2A).

### Relationship between HDI-transfected enhancer reporter activity and TCPOBOP-responsive epigenetic features

The above findings demonstrate that condition-specific liver enhancer activity can be assayed over a large dynamic range by HDI delivery of reporter plasmids. Further, the DHS sequences that showed large TCPOBOP-induced increases in enhancer activity (DHS #50358, DHS #29774) showed increased chromatin opening in mouse liver following TCPOBOP treatment [58]. Both DHS also showed significant increases in flanking enhancer marks (H3K27ac, but not H3K4me1), binding of the nuclear receptor CAR, and target gene expression (Table S2A) in mouse liver. In the case of DHS #29774, HDI reporter activity was increased only 4.9-fold following TCPOBOP treatment, but from a comparatively high basal level, whereas the presumed target gene, *Cyp2c55*, showed a much greater, 182-fold increase in expression after TCPOBOP treatment. This discrepancy in *Cyp2c55*-associated enhancer activity vs gene responses suggests that interactions with one or more of the 3 other nearby TCPOBOP-opened DHS sequences may additionally be required to recapitulate the robust induction exhibited by the endogenous gene. In contrast, DHS #50358 displayed very low basal reporter activity but with a much stronger (245-fold) increase following TCPOBOP treatment. This increase is sufficient to account for the dramatic induction of the adjacent *Cyp2b10* (125-fold increase) and the divergently transcribed lncRNA gene, *lnc5998* (375-fold increase). Finally, results with DHS #50363 and DHS #5898 (low basal reporter activity and little or no increase with TCPOBOP treatment) exemplify that only a subset of TCPOBOP-activated enhancer DHS are functional for enhancer activity in vivo using HDI, despite significant TCPOBOP-stimulated increases in chromatin opening, H3K27ac enhancer marks and/or induced CAR binding at the corresponding endogenous liver gene sequences (Table S2A).

### Optimized time point for HDI reporter assay

HDI-transfected plasmid DNA levels are initially very high in mouse liver, but rapidly drop off after 1 d and thereafter remain stable for many weeks [51, 69]. Renilla luciferase reporter activity (pGYL18) showed the same rapid drop-off, decreasing by 9 to 10-fold from 20 h to 7 d after HDI (Fig. S1A). Even more dramatic decreases in firefly luciferase activity were seen from 20 h to 7 d for the *Alb* promotor empty vector and when transcription from that same promoter was driven by an *Alb* enhancer (pGYL17; decreases of 212-fold and 30-fold, respectively; Fig. S1A). These decreases are likely due to the elimination of free plasmid DNA from the liver and/or changes in plasmid chromatinization in vivo over time [70]. Although absolute reporter activity was higher after 20 h, the much higher background (empty plasmid) reporter activity at 20 h as compared to 7 d resulted in a dramatic, 11.6-fold increase in signal to noise ratio for the reporter plasmid after 7 d as compared to 20 h (63.6 vs 5.46; Fig. 1C). HDI-transfected plasmid DNA remaining in the nucleus after 7 d is stably associated with liver cells and shows long term expression [59, 71]. This can be explained by the transfected episomal DNA adopting a nucleosomal structure, as was shown by immunoprecipitation [72], in situ hybridization [73] and selective hybridization followed by sequencing [74]. On this basis, we selected 7 d after HDI as the time point for determination of liver enhancer activity.

### STARR-seq plasmids do not induce strong type-I interferon responses in mouse liver following HDI

Naked bacterial plasmid DNA elicits a strong innate immune response in cultured cells, most notably activation of type-I interferon response genes, which is a major confounding factor that impacts the reliability of enhancer activity assays such as STARR-seq [25]. To ascertain whether this limitation applies to HDI delivery of pPromALB-based STARR-seq reporter plasmids to mouse liver, we examined the effects of STARR-seq plasmid transfection by HDI on a panel of type-I interferon response genes previously shown to be strongly induced by STARR-seq plasmid transfection in cell lines [25]. qPCR analysis of RNA extracted from the HDI-transfected livers revealed minimal type-I interferon gene responses after 7 d; moreover, liver expression of the established type-I interferon response genes *Cxcl10*, *Ifi27*, *Ifit1* and *Irf7* was significantly decreased after 7 d when compared to the expression level 20 h after HDI (Fig. 1D). Furthermore, no significant differences in the liver interferon response were seen following HDI transfection of STARR-seq plasmids incorporating the *Alb* promoter as compared to the ORI or SCP1 promoters (Fig. S2), both of which elicit strong type-I interferon responses in multiple cultured cell lines [25]. In addition, liver histology was normal after 7d (Fig. S3), consistent with the resolution of hepatocyte swelling and other liver morphological changes and tissue damage within 6-24 h after HDI [75]. We conclude that a 7 d waiting period following HDI is sufficient to insure a low baseline level of type-I interferon response and resolution of any initial, associated liver pathology, and that liver HDI reporter activity can be determined directly, without the need to treat with pharmaceutical inhibitors to suppress the innate immune response [25].

### Focused (100-plex) HDI-STARR-seq library

We tested the ability of HDI-STARR-seq to assay open chromatin regions in a multiplexed manner by generating a STARR-seq plasmid library comprised of PCR-amplified genomic DNA fragments targeting 100 mouse liver DHS sequences (Fig. 2A, Table S1C, Table S1D). The DHS sequences examined ranged in size from 297-2,038 nt (mean = 686 nt; median = 556 nt) and included 36 genomic regions whose chromatin accessibility is either induced or repressed following exposure to TCPOBOP [58] and 44 genomic regions whose chromatin accessibility differs significantly between male and female liver and may contribute to sex biased regulation of liver gene expression [40]. PCR-amplified genomic DNA fragments longer than 600 bp were lightly digested with DNase-I to generate a pool of shorter DNA fragments (size range 100-500 bp; Fig. S4A) to facilitate finer mapping of active subregions within a DHS region. The DNA fragments obtained were mixed with PCR amplicons shorter than 600 bp and then ligated *en masse* to SalI/AgeI-digested pTYC10 (pPromALB_STARR-seq; Table S1E). The final pool of PCR fragments ranged in size 100-600 bp (Fig. S4B).

The resultant STARR-seq library was successfully delivered to individual male and female mouse livers by HDI, as verified for each liver using a co-transfected Renilla luciferase reporter plasmid (Fig. S1B). Livers were collected 7 d later, in some cases from mice treated with TCPOBOP (or vehicle control) 24 h prior to tissue collection to activate endogenous, CAR-dependent enhancer DHS [58]. Three types of libraries were prepared for Illumina sequencing: the original plasmid library used for HDI delivery (Input plasmid library), plasmid DNA recovered from individual HDI-transfected livers (HDI liver DNA library), and poly-adenylated RNA reporter sequences extracted from the HDI livers and then reverse transcribed to cDNA (STARR-seq reporter RNA library) (Fig. 2B). MACS2 peak analysis of mapped sequence reads combined across all biological replicate libraries identified 1,673 peak regions genome-wide, of which the 100 targeted PCR-amplified DHS sequences represented 90-93% of the STARR-seq RNA library sequence reads mapping to the 1,673 peak regions (Table S3A, Library STARR_TYC3/7).

HDI delivery of the focused STARR-seq library was highly reproducible between individual mice, as seen by comparing normalized read counts across the 100 DHS regions for the original input plasmid library to those for libraries prepared from plasmid DNA extracted from male, female and TCPOBOP-treated male livers (r^2^ = 0.968-0.978, Fig. 3A). Importantly, the HDI-STARR-seq RNA activity profiles of the individual DHS regions, normalized by the plasmid sequence read counts of each region, clustered by mouse group (Fig. 3B).

**Fig. 3.**
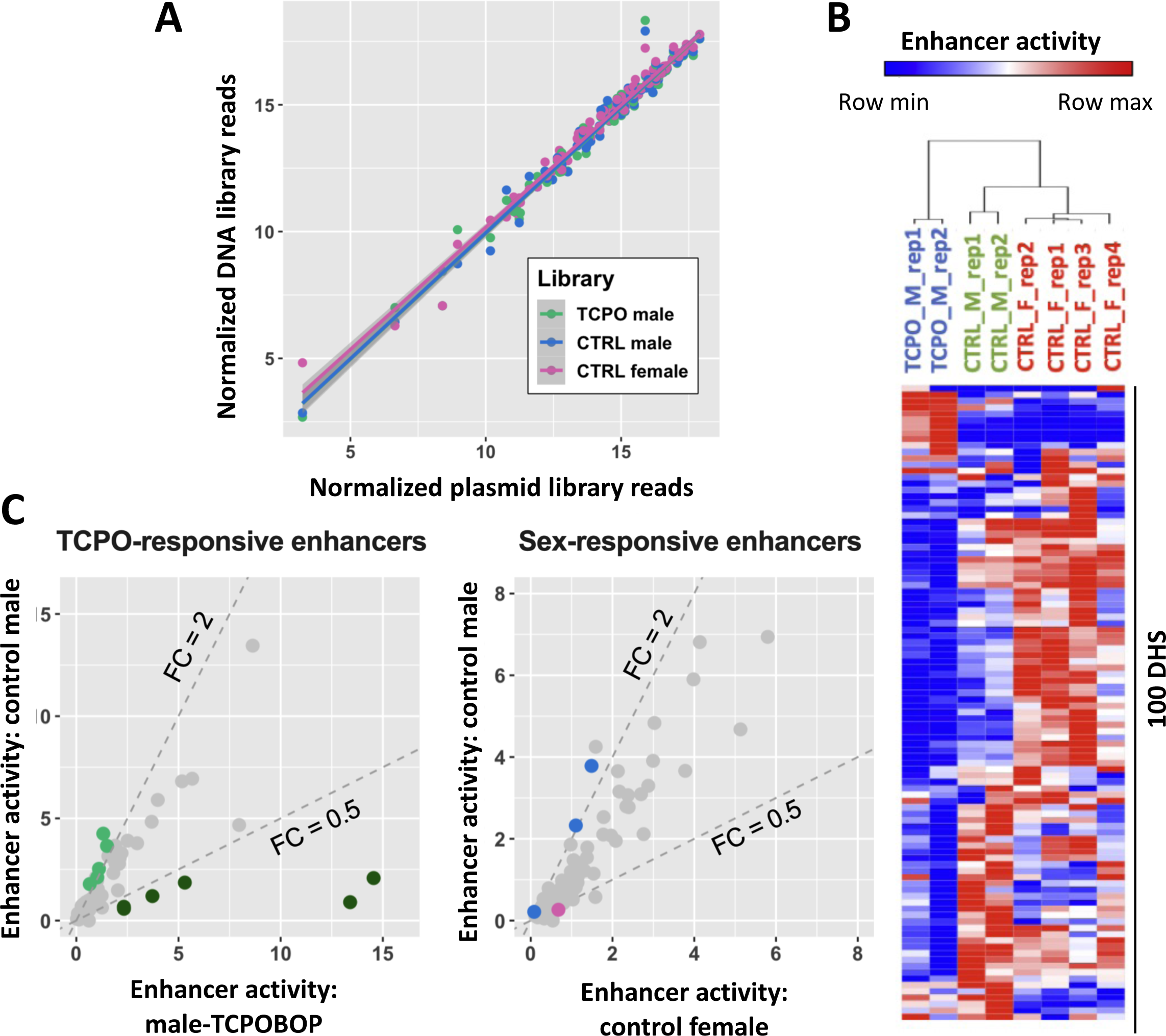
HDI-STARR-seq using focused library comprised of 100 DHS. (A) Consistency on log2 normalized sequence reads across 100 targeted DHS regions between input plasmid library (X-axis) and DNA library extracted from one mouse liver for each of the 3 indicated biological conditions (liver with highest Renilla luciferase activity; Fig. S1B), with r^2^ = 0.968 (TCPOBOP-treated male, *green*), 0.975 (control male, *blue*) and 0.978 (control female, *pink*), respectively. (B) Heatmap showing hierarchical clustering (Morpheus; https://software.broadinstitute.org/morpheus) of RNA activity profiles across biological replicate livers for each experimental condition. Blue represents the minimal enhancer activity among 8 livers, and red represents the maximal enhancer activity among 8 livers. (C) Pairwise analysis of enhancer activities for the 100 DHS identifies condition-specific enhancers for TCPOBOP responsiveness (left) and for sex specificity (right), with colored dots marking enhancers that show >2-fold difference in activity between conditions at p < 0.05.

Moreover, as discussed below, comparison of HDI-STARR-seq activity profiles between mouse groups (males, females and TCPOBOP-treated males) identified many condition-dependent enhancers (Fig. 2C, Fig. 3C, Fig. S5, Table S3B), supporting the utility of HDI-STARR-seq for condition-specific enhancer characterization.

### TCPOBOP-responsive enhancers: relationship between HDI-STARR-seq activity and liver epigenetic features

26 of the 100 DHS regions tested showed a significant difference in liver HDI-STARR-seq enhancer activity (p < 0.05, unpaired Student *t*-test) 24 h after TCPOBOP treatment (Table S3B, column AX; Fig. 4A). Enhancer activity was increased by TCPOBOP at 7 of the 26 DHS regions, including both of the TCPOBOP-activated enhancers identified by HDI luciferase assay (Fig. 1B, Fig. 2C, Fig. S5A, Fig. S5B), and activity was decreased at the other 19 DHS regions (Table S3B, columns AT-AX). All 7 DHS where TCPOBOP stimulated enhancer activity undergo robust chromatin opening in mouse liver following TCPOBOP treatment, with 5 of the 7 DHS regions also showing an increase in the activating histone mark H3K27ac (Table S3C, columns U and AK) [36, 58]. Thus, these DHS harbor regulatory elements that are activated when TCPOBOP-bound CAR increases their accessibility in mouse liver chromatin, and when tested as episomal sequences by HDI-STARR-seq. None of the 7 TCPOBOP-activated enhancers contained endogenous repressive chromatin marks (i.e., H3K27me3; Table S3C, column AT). In contrast, 13 of the 19 DHS regions whose HDI-STARR-seq enhancer activity was decreased by TCPOBOP were static, i.e., TCPOBOP exposure did not induce endogenous chromatin opening or chromatin closing in mouse liver, and activating histone marks were either absent or unchanged following TCPOBOP treatment at 16 of the 19 DHS regions (Table S3C, columns U, AK-AM). Of note, the repressive mark H3K27me3 was absent from the 19 repressed DHS regions. Finally, TCPOBOP had no significant effect on the HDI-STARR-seq activity of 55 DHS regions whose chromatin accessibility was unchanged by TCPOBOP (static DHS) or on the activity of 17 other DHS where TCPOBOP induced chromatin opening (Table S3C, columns O and U).

**Fig. 4.**
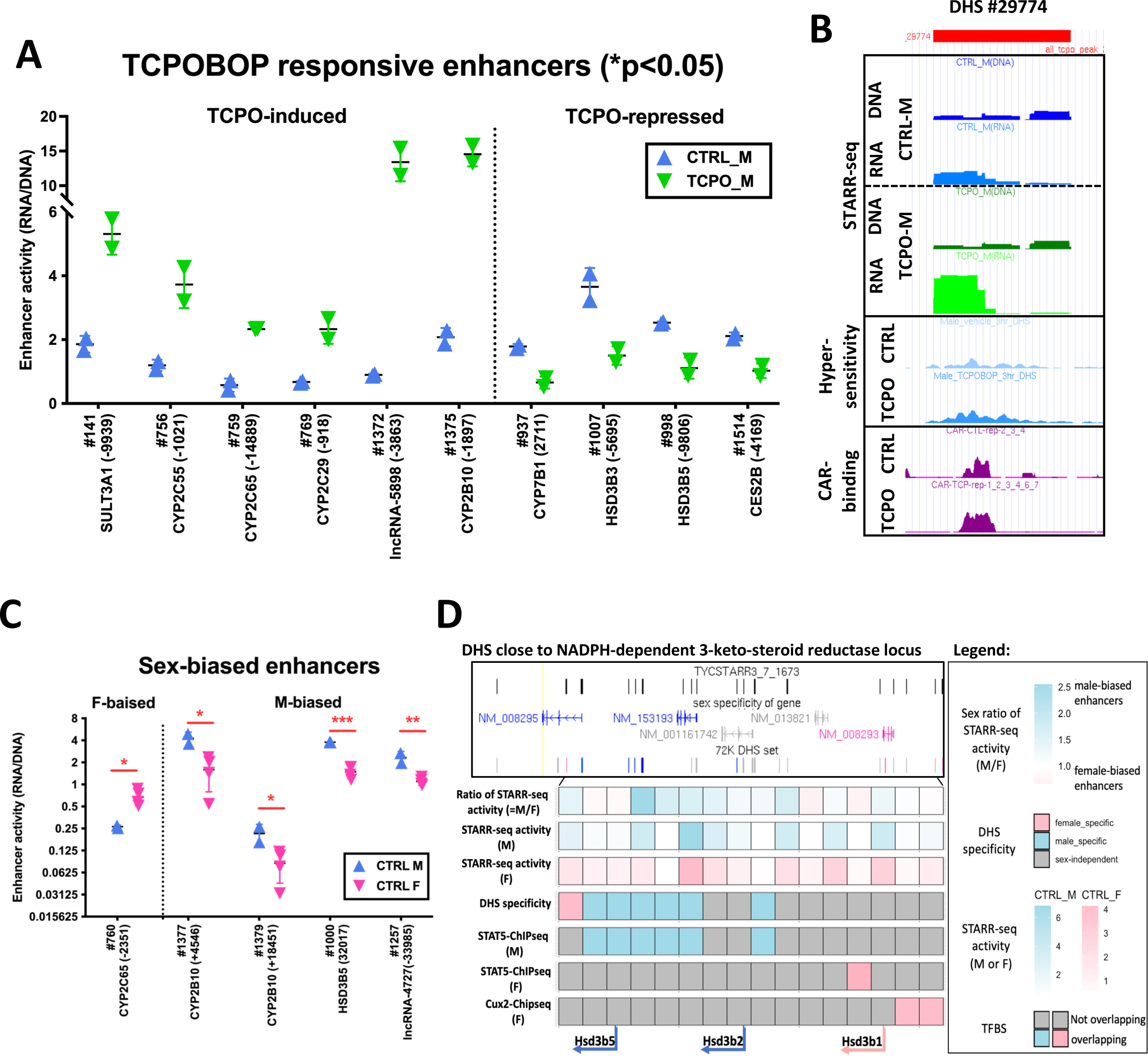
Condition-specific enhancers identified by HDI-STARR-seq: xenobiotic response and sex specificity. (A) HDI-STARR-seq enhancer activity for 10 focused library DHS showing significantly differential reporter activity, either induced or repressed by TCPOBOP treatment (*, p<0.05 by *t*-test, |fold-change|>2). In addition, 16 other DHS responded to TCPOBOP significantly, but < 2-fold (Table S3B, column AX). The closest protein coding gene is indicated below the x-axis, and the nt distance from the gene TSS to the DHS is in the parenthesis (positive value, DHS downstream of the TSS; negative value, DHS upstream of the TSS). Data shown are mean activity values +/- SEM for n=2 livers per condition. (B) UCSC browser tracing showing STARR-seq DNA and RNA activity for DHS # 29774 on chr19, upstream of Cyp2c55, in control male liver (top 2 tracks) and in TCPOBOP-treated male liver (next 2 tracks), followed by tracks showing DNase hypersensitivity and CAR binding activity. (C) HDI-STARR-seq enhancer activity for 5 focused library DHS showing sex-differential enhancer activity, as in panel A. Data are shown for n=2 for male and n=4 female livers, with *, p<0.05; **, p<0.01; and ***, p<0.001. (D) STARR-seq enhancer activity across a genomic region with sex-biased *Hsd3b* family genes. 16 DHS are located in the same TAD regions as three sex-biased genes, including male-biased genes (*Hsd3b5*, *Hsd3b2*) and a female-biased gene (*Hsd3b1*). The indicated genomic region is subdivided into 16 sequential subregions (16 horizontal boxes); the labels at the left of each row indicate the dataset displayed in that row, i.e., STARR-seq enhancer activities in male versus female liver, individual STARR-seq enhancer activities, DHS sex specificity, STAT5 ChIP-seq peaks in male liver and in female liver, and female-specific Cux2 ChIP-seq peaks in female liver, with signal intensities indicated in the legend.

**Fig. 5.**
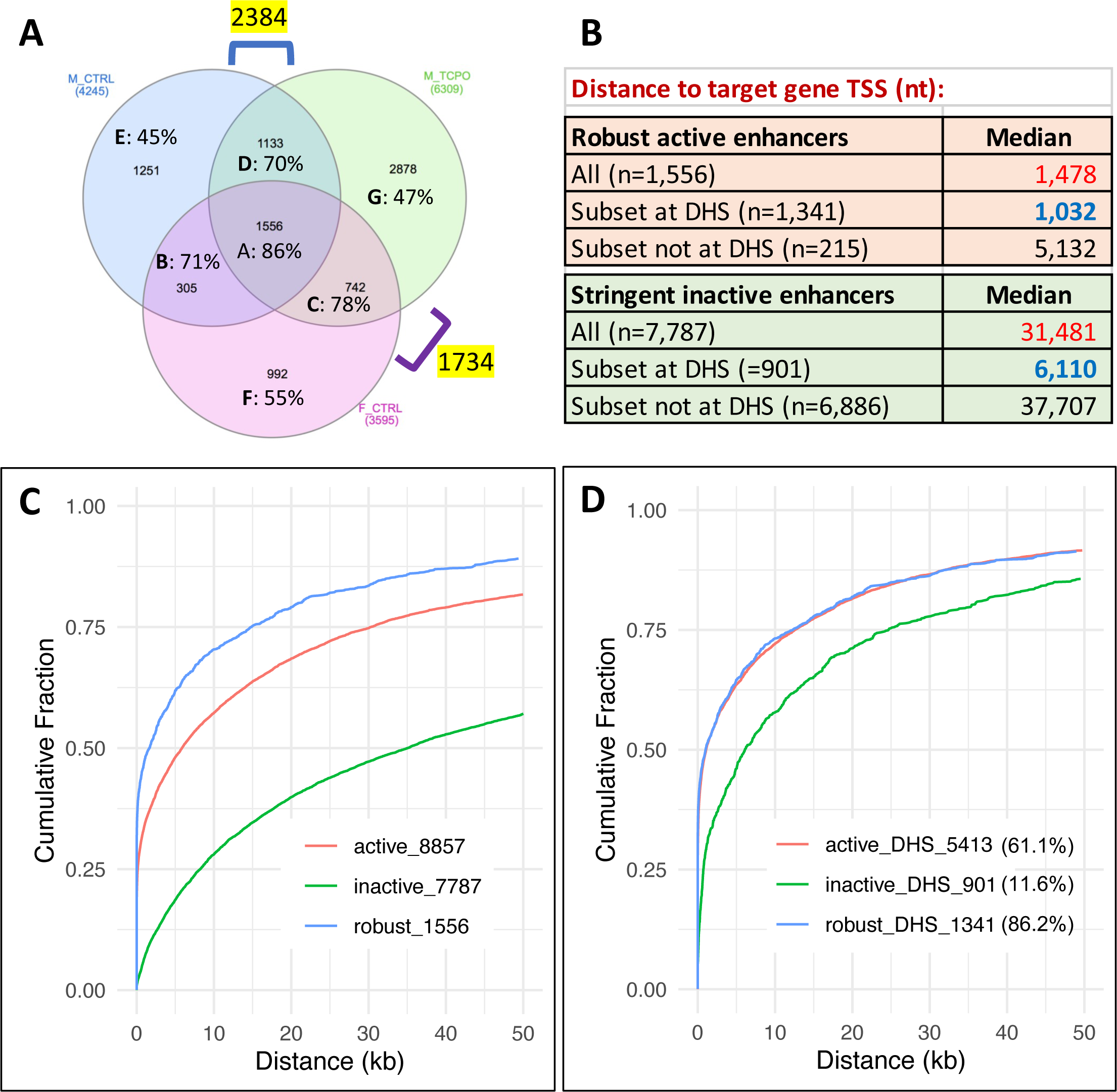
HDI-STARR-seq active vs stringently inactive enhancers: differential proximity to TSS: (A) Venn diagram showing numbers of active enhancers identified using global HDI-STARR-seq library in male (control) liver, female (control) liver, and TCPOBOP-treated male liver. A total of 1,556 enhancers (Group A) show activity under all 3 biological conditions and are designated robust active enhancers. A total of 4,245 enhancers were active in male liver, 6,309 in TCPOBOP-treated male liver and 3,595 in female liver, as indicated outside of each circle. Yellow highlight: numbers of enhancers active in male but not female liver (2,384), and in female but not male liver (1,734). Numbers in parenthesis indicate the percentage of genomic regions in each set that overlaps a liver DHS. Also see Fig. S6C and Table S4. (B) Median distance of each enhancer set to the nearest gene TSS in the same TAD. Also see Table S4C, Table S4D. (C, D) Plots showing the cumulative percentage of enhancers in each indicated enhancer set as a function of distance from either end of an HDI-STARR-seq enhancer to the nearest gene TSS (Y-axis). Analysis was performed in R Studio and plotted using ggplot2. The subsets of each enhancer set in panel C that overlap a DHS are presented in panel D.

The 7 TCPOBOP-activated enhancers mapped to a total of 6 TCPOBOP-induced genes: *Cyp2b10* and the divergently transcribed *lnc5998* (*nc_inter_c7_5998)* [58], *Cyp2c29*, *Cyp2c55*, *Cyp2c65* and *Sult3a1*, with distances to the gene TSS ranging from 0.9 kb to 14.8 kb (Table S3C, columns V-AB). Of note, the 5 activated enhancers that showed an increase in endogenous H3K27ac marks were proximal (< 5 kb) to their TCPOBOP-induced gene targets, and were bound by CAR (e.g., Fig. 4B) and its heterodimerization partner RXR (Table S3C, columns AW, BD, BE), suggesting direct CAR binding is a key feature of their TCPOBOP-induced enhancer activity. In contrast, 2 of the TCPOBOP-activated enhancer DHS (Focused peaks 141 and 759, Fig. 4A) were devoid of CAR binding and of activating histone marks, implicating CAR-independent mechanisms in TCPOBOP-stimulated endogenous chromatin opening at these sites, and in TCPOBOP induction of HDI-STARR-seq enhancer activity. None the 19 DHS regions where TCPOBOP repressed HDI-STARR-seq reporter activity mapped to TCPOBOP-repressed genes and only 2 of those 19 DHS harbor CAR-binding sites, consistent with indirect mechanisms regulating enhancer repression (Table S3C, columns AD, AW, BA, BD). Unexpectedly, 9 of the 19 TCPOBOP-repressed enhancers mapped to a TCPOBOP-induced gene, suggesting they may act to moderate TCPOBOP gene induction (Table S3C, column AD).

### Sex-biased enhancer activity

Comparison of HDI-STARR-seq enhancer activity between male and female livers identified 13 male-biased and 16 female-biased enhancers at P < 0.05 (Table S3B, column AX), of which 5 showed >2-fold difference in enhancer activity between the sexes (stringent sex bias; Fig. 4C). One of the stringent male-biased enhancers, Focus_MACS2_Peak_1000, mapped to *Hsd3b5*, which shows male-biased expression in mouse liver (Table S3D). This enhancer is characterized by epigenetic regulatory features consistent with its role in growth hormone-regulated, male-biased gene expression [40, 76]. Thus, it corresponds to a liver DHS that binds the growth hormone-activated TF STAT5 in a male-biased manner, and exhibits chromatin accessibility that is significantly greater in male than female liver and is significantly decreased in male mice infused with growth hormone continuously for 7-10 days, which mimics the female plasma growth hormone profile and leads to closure of many male-biased DHS and loss of male-biased gene expression, including that of *Hsd3b5* [77]. Of note, Focus_MACS2_Peak_1000 maps to a TAD that contains both male-biased genes (*Hsd3b2*, *Hsd3b5*) and female-biased genes (*Hsd3b1*). The 5’ genomic segment, which includes *Hsd3b2* and *Hsd3b5*, harbors genomic sequences with male-biased chromatin accessibility (DHS), male-biased STAT5 binding sites and regions with male-biased HDI-STARR-seq enhancer activity, while the 3’ genomic segment, which includes *Hsd3b1*, contains female-biased DHS, female-biased binding sites for STAT5 and for the female-specific growth hormone-regulated TF Cux2, and female-biased HDI-STARR-seq enhancer activity (Fig. 4D, Fig. S5D, Table S3E).

The stringent female-biased enhancer identified, Focus_MACS2_Peak_760, maps to a TAD with 5 female-biased genes from the *Cyp2c* subfamily (*Cyp2c37*-*Cyp2c40*, *Cyp2c69*), although the closest gene, *Cyp2c65*, does not show sex-biased expression. However, not all genomic regions that showed a sex bias in HDI-STARR-seq enhancer activity mapped to genes showing the same sex bias in gene expression (e.g., male-biased HDI-STARR-seq activity at Focus_MACS2_Peak_1377 and Focus_MACS2_Peak_1379, which both map to the female-biased gene *Cyp2b10*). One possible explanation is that HDI-STARR-seq measures the intrinsic transcriptional capability of a single enhancer, while sex-biased gene expression may reflect the functional integration of multiple sex-biased enhancers and repressors.

### Impact of promoter choice on liver HDI-STARR-seq enhancer activity

Next, we investigated the effectiveness of HDI-STARR-seq for interrogating liver DHS regions on a global scale, as well as its ability to provide finer resolution within active DHS regions. Genomic DNA fragments released by DNase-I digestion of mouse liver nuclei (i.e., open chromatin regions) were extracted from both untreated and TCPOBOP-treated mouse livers and pooled. The fragments were cloned in an orientation-independent manner into pPromALB-STARR-seq plasmid (library STARR-TYC6), which contains the same minimal *Alb* promoter used in our HDI luciferase assays (Fig. 1B). Cloning was performed using multiple, parallel PCR reactions in an effort to minimize PCR-dependent loss of library complexity. In parallel, the same batch of DNase-I released open chromatin fragments was cloned into two other STARR-seq libraries that differed only in the promoter sequences used to drive transcriptional initiation of reporter activity: Super Core 1 (SCP1), a synthetic promoter (library STARR-TYC4) and ORI, a bacterial origin promoter (library STARR-TYC5), both commonly used for STARR-seq [25]. Each library was delivered to mouse liver by HDI, followed by extraction of plasmid DNA and reporter RNA using the same amount of tissue from each liver. Although the recovery of plasmid DNA from the HDI-transfected livers was similar for all 3 STARR-seq libraries, the yield of PCR-amplified reporter RNA from the ORI-STARR-seq and SCP1-STARR-seq libraries was 5-10-fold lower than for the pPromALB-STARR-seq library. Furthermore, after sequencing, the total mapped RNA read counts were 50 to 150-fold lower for the ORI and SCP1 reporter RNA libraries (mean mapped reads: 0.57, 1.9 and 89.5 million mapped fragments for SCP1, ORI and pPromALB STARR-seq RNA libraries, respectively; c.f., 41-56 million mapped recovered plasmid DNA sequence reads, respectively; Table S3A). Thus, the SCP1 and ORI promoters do not support HDI-STARR-seq reporter activity in mouse liver.

### Global HDI-STARR-seq library: evaluation of enhancer activity under diverse biological conditions

HDI-STARR-seq analysis of the global liver DHS plasmid library, pPromALB-STARR-seq, was performed in both male and female mice, and in male mice given TCPOBOP 24 h prior to liver collection 7 d after HDI. Illumina sequencing of the input plasmid library, followed by MACS2 analysis, identified 50,332 qualified autosomal peaks (minimum of 40 Illumina sequence reads per 10 million mapped reads), with a width of 258 +/- 97 nt, mean +/- SD (Table S4A). 14,330 (28%) of the 50,322 peaks overlap open chromatin regions (i.e., DHS) identified previously in mouse liver [37, 58] (Table S4A). The presence of many non-DHS peak regions in the global library is consistent with our finding that ∼15-25% of DNase-released fragments used to prepare the global library localize to DHS peak regions [58].

We set a threshold for identification of active enhancers, as follows. First, we observed that genomic regions with HDI-STARR-seq library enhancer activity confirmed (at a threshold of > 20 normalized RNA reads/10 million mapped reads) in 2 or more biological replicate livers showed a liver number-dependent pattern of increasingly high STARR-seq library sequence representation (Fig. S6A). We therefore applied stringent criteria to define a genomic region as an active enhancer, namely, the region met our threshold for HDI-STARR-seq activity in at least 3 individual livers for a given biological condition (Fig. S6B). Overall, 8,857 enhancers showed consistent STARR-seq activity under at least one of the 3 biological conditions tested (Table S4C); moreover, 1,556 of the 8,857 enhancers showed enhancer activity under all 3 biological conditions (female, male and TCPOBOP-treated male livers), and were designated robust HDI-STARR-seq enhancers (Fig. 5A). We also identified 7,787 stringently inactive enhancers, defined as genomic regions within the set of 50,332 qualified genomic regions that gave ≤ 20 reporter RNA library reads per 10 million mapped reads for each of the 11 livers examined (i.e., all 4 untreated male livers, all 4 TCPOBOP-treated male livers and all 3 untreated female livers) (Table S4C). These stringently inactive enhancers were used as a background set for enrichment analysis (see below).

### Enrichment for open chromatin regions

The active enhancers were highly enriched for mouse liver open chromatin regions: 86% of the 1,556 robust enhancers overlapped a liver DHS region identified previously [37, 58], whereas only 11.6% of the 7,787 stringently inactive enhancers overlapped with a liver DHS region (ES = 49.7, p< E-05). These percentages (86%, 11.6%) can be compared to expected values of 37% and 28% overlap with a liver DHS for the robust active and for the stringently inactive enhancers, respectively, taking into account the moderate dependence of DHS frequency on representation within the global STARR-seq plasmid library (Table S4D). Moreover, 5,413 (38%) of the 14,330 DHS regions represented in the global HDI-STARR-seq library showed enhancer activity under at least one biological condition, as compared to only 3,444 (9.6%) of 35,992 global library regions not mapping to a DHS (ES = 5.8; Table S4C). The percentage of sites overlapping open chromatin regions decreased to 71-78% when we considered active enhancers identified in only 2 of the 3 biological conditions (Venn regions B-D, Fig. 5A) and it decreased further, to 45-55%, for active enhancers unique to one biological condition (Venn regions E-G) (Fig. S6C). This pattern suggests a higher false positive rate for HDI-STARR-seq enhancers identified under only one biological condition, which may reflect noise introduced for genomic regions with lower representation in the STARR-seq plasmid library (Fig. S6D).

### Enrichment for TSS proximal sequences

Mapping to target genes (closest TSS) revealed that 52% of robust active enhancers were TSS proximal (< 2 kb), as compared to only 11% for stringently inactive enhancers (ES = 8.7-fold, p=2.4E-258; Table S4D). This contrasts with STARR-seq active enhancers identified in transfected GM12878 cells, a majority (77%) of which are TSS distal (> 2 kb) [78]. Overall, the enhancer to nearest TSS median distance was >20-fold closer for the set of robust active enhancers (1,478 nt vs. 31,481 nt for stringently inactive enhancers) (Fig. 5B, Fig. 5C). The same trend was seen for the STARR-seq enhancer subsets that overlap a liver DHS, which are closer to gene TSSs than active enhancers not at a DHS: 1,032 nt (median) distance to nearest TSS for robust active enhancers at a DHS vs. 6,110 nt for stringently inactive enhancers at a DHS (Fig. 5D, Table S4D). We conclude that DHS sequences proximal to gene TSSs are more likely to show activity HDI-STARR-seq activity than gene TSS distal DHS sequences.

### Epigenetic features of active enhancers

We investigated whether the set of 1,556 robust active enhancers was enriched for liver genomic regions with activating or repressive histone marks (Fig. 6A), or for particular chromatin states (Fig. 6B), based on extensive published epigenetic datasets for mouse liver [40]. Strikingly, the robust active enhancers showed strong enrichments for histone marks associated with active enhancers (ES = 31.7 for H3K27ac, ES = 11.4 for H3K4me1) and active promoters (ES = 15.5 for H3K4me3), as well as significant depletion of repressive histone marks (ES = 0.16 to 0.25, for H3K27me3 and H3K9me3) when compared to the background set comprised of 7,787 stringently inactive enhancers. The transcribed region mark H3K36me3 was also significantly depleted (ES = 0.41) (Fig. 6A). Enrichments for activating histone marks were also high, albeit somewhat lower, when we considered the full set of 8,857 active enhancers, but were much lower for 33,688 other genomic regions whose HDI-STARR-seq activity did not meet our threshold for designation as active enhancers (Fig. 6A). Consistent with these findings, the set of robust active enhancers was strongly enriched for chromatin states E7 and E5, and to a lesser extent state E6; significant depletion of several inactive and transcribed chromatin states was also observed (Fig. 6B). State E7 is characterized by open chromatin combined with the promoter mark H3K4me3 and the enhancer mark H3K27ac; its strong enrichment is consistent with our finding, above, that many robust active enhancers are close to gene TSSs. E5 and E6 are enhancer states rich in open chromatin regions, either without or with enhancer histone marks H3K4me1 and H3K27ac. Enrichments for other active enhancer subsets are shown in Fig. S7 and Fig. S8, with full results in Table S4E.

**Fig. 6.**
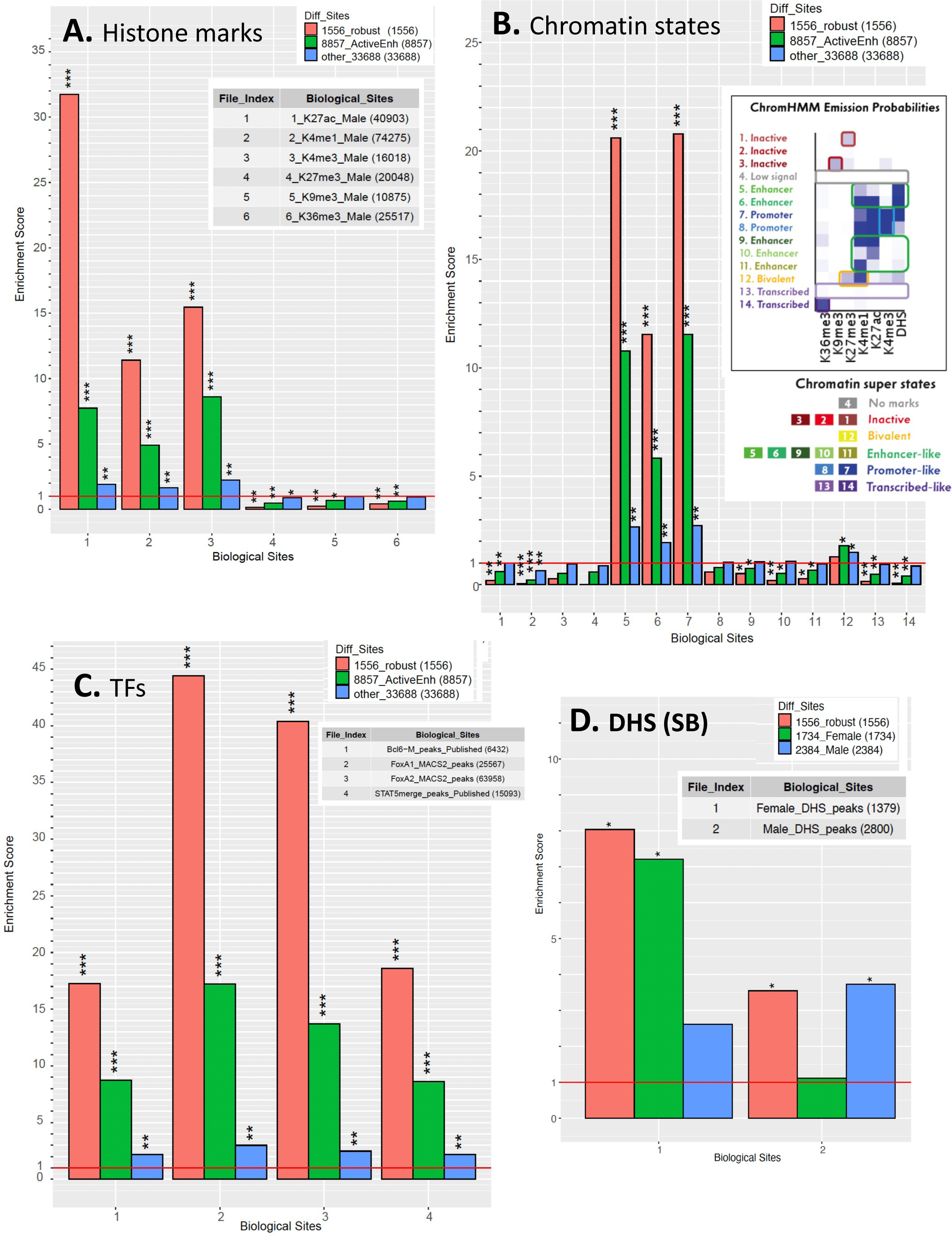
HDI-STARR-seq active enhancers: enrichment for epigenetic features and TF binding in mouse liver. Enrichment scores for the overlap between active enhancers identified by HDI-STARR-seq (Fig. 5A) and the sets of epigenetic and TF binding features indicated in each panel, with the number of features shown in parenthesis. Enrichments in A, B, and C were determined for 3 active enhancers sets: the set of 1,556 robust active enhancers (*salmon*), the full set of 8,857 active enhancers (*green*), and the set of 33,688 other enhancers that show enhancer activity in at least one of 11 of the individual livers tested (*blue*) (see Fig. S6C). Enrichments were computed relative to a background set of 7,787 stringently inactive genomic regions. Shown are enrichments for: (A) Histone H3 chromatin marks identified by ChIP-seq in male mouse liver; (B) occurrence of each of 14 chromatin states, determined for male liver by ChromHMM analysis, with the emission probabilities describing each state shown in the inset; (C) TF binding sites for the 4 indicated TFs. (D) shows sex-specific DHS for active enhancers identified in male and female livers (marked outside of Venn diagram in Fig. 5A). The significance of enrichment of each active enhancer set was determined by Fisher’s exact test: *, P<0.001; **, P<1e-10; ***, P<1e-100. Full data are shown in Table S4E.

We also identified active enhancer subsets that showed HDI-STARR-seq activity in male but not female liver (n=2,384, Venn regions D + E) or in female but not male liver (n=1,734, Venn regions C +F) (Fig. 5A). Importantly, the male-biased enhancers were significantly enriched for mouse chromatin regions showing greater chromatin accessibility in male liver than in female liver, and correspondingly, the female-biased enhancers were significantly enriched for mouse chromatin regions showing greater chromatin accessibility in female liver than in male liver (Fig. 6D). Thus, the sex-dependence of enhancer activity matches the sex-dependent pattern of endogenous chromatin accessibility seen in mouse liver, which itself is linked to the sex bias of gene expression [40]. Moreover, the full set of robust active enhancers showed strong enrichment for liver binding sites for several TFs regulating transcription of sex-dependent genes [63, 79] (Fig. 6C).

Finally, we investigated the arrangement of enhancer features flanking the STARR-seq active enhancers by considering three sequential 400 bp sub-regions centered on each HDI-STARR-seq active region (Fig. S9A). Whereas the central sub-region showed strong enrichment for chromatin states E5 and E7, and to a lesser extent state E6, the flanking regions were most highly enriched for the active promoter state E7, consistent with the close proximity of many active enhancers to gene TSSs indicated above, followed by enhancer states E5 and E6, and to a lesser extent promoter state E8 and enhancer state E9. States E8 and E9 are populated by active histone marks but are relatively depleted of open chromatin regions, as is to be expected for sequences that flank enhancer DHS regions (Fig. S9B-S9G). Taken together, the strong associations between HDI-STARR-seq enhancer activity and endogenous epigenetic marks and chromatin states, as well as liver transcription binding patterns, indicate that the HDI-STARR-seq reporter assay recapitulates the enhancer activity, and perhaps also the chromatin state, of the corresponding endogenous genomic sequences in mouse liver.

## Discussion

STARR-seq and other MPRAs enable enhancer activity to be assayed in parallel for thousands of DNA sequences and is commonly employed in cell culture studies using transfected reporter plasmids. Major limitations include the lack of a native chromatin structure for the transfected plasmid DNA, and the absence of the three-dimensional, multi-cellular environment of a complex tissue, where factors such as the extracellular matrix, cell-cell interactions and exogenous hormonal stimuli impact the activities of thousands of enhancers. To address these issues, we build upon STARR-seq [23] to develop HDI-STARR-seq, which combines STARR-seq plasmid library delivery to the liver, by HDI, with the introduction of a minimal *Alb* promoter, which we show is absolutely essential to measure significant STARR-seq enhancer activity in mouse liver in vivo. We established the utility of this method for assaying enhancer activity in liver in vivo using a focused HDI-STARR-seq library and validated select results in vivo using luciferase reporter assays.

Comparisons of HDI-STARR-seq enhancer activity in male vs. female mouse liver, and in livers of male mice treated with TCPOBOP, a foreign chemical agonist ligand of the liver TF CAR (*Nr1i3*), identified condition-dependent enhancers associated with condition-specific gene expression. Importantly, we established that HDI-STARR-seq does not activate a strong plasmid DNA-induced type-I interferon immune response, which confounds STARR-seq enhancer analysis in transfected cell models [25]. Further, we demonstrated that HDI-STARR-seq can readily be scaled for global interrogation of 50,000 genomic regions, which were cloned into the HDI-STARR-seq reporter plasmid as DNA fragments released by light DNase-I digestion of purified liver nuclei (open chromatin regions). Thousands of active liver enhancers were identified and shown to be significantly enriched for genomic regions close to gene TSSs with activating histone marks (H3K27ac, H3K4me1, H3K4me3) and depleted of repressive (H3K27me3, H3K9me3) and transcribed region marks (H3K36me3) in mouse liver. Thus, HDI-STARR-seq can readily be implemented on a global scale to characterize condition-specific regulatory sequences that are functionally active in liver in vivo under a range of biological conditions.

### HDI delivery of STARR-seq reporter plasmids to mouse liver

HDI is an effective way to transfect liver cells with plasmid DNA in an intact mouse without the need to prepare chromatinized plasmids or viral vectors. HDI is highly specific for plasmid delivery to the liver [51] and has been implemented in many species, including rat, rabbit and porcine liver [80–82]. Transfection efficiencies in mice range from 5% to 40% of liver cells [83], depending on injection conditions. HDI requires rapid tail vein injection of plasmid DNA in a large volume, which induces large increases in intravascular pressure and membrane permeability and is accompanied by a transient increase in intrahepatic vesicles and hepatocyte swelling, primarily in the pericentral region, where transgene expression dominates [75, 84]. Liver non-parenchymal cells are also subjected to these hydrodynamic forces, but their expression of HDI-delivered episomal plasmids is low [84]. Plasmid retention and transgene expression persist for many weeks [59, 69], making HDI a useful tool for studying liver promoters and enhancers [38, 85]. Importantly, we found that delaying liver collection until 7 d after HDI, when the large excess of free plasmid has been eliminated from the liver [69], significantly increased the signal-to-noise ratio for liver enhancer reporter activity, which may help minimize false positives linked to the initially very high level of transgene expression seen following HDI [59, 69]. Co-injection of a Renilla luciferase reporter allowed us to identify and exclude livers with low transfection efficiency. The large interindividual variation in the efficiency of HDI (Fig. S1B) is likely a consequence of the large impact that small variations in the total actual time required for each injection have on the level of transgene expression [51].

### HDI-STARR-seq does not induce a sustained type-I interferon response

Introduction of naked plasmid DNA in STARR-seq-based transfection experiments elicits a strong type-I interferon innate immune response [25]. This response, which can also occur when plasmid DNA is transfected into mouse liver [86], is a serious confounding factor for STARR-seq, as it produces large numbers of enhancer false positives and false negatives [25], raising concerns about the reliability of enhancer activities determined by STARR-seq, even when kinase inhibitors are added to block type-I interferon induction [87]. In contrast, using HDI-STARR-seq, we found little or no sustained type-I interferon response when naked STARR-seq plasmid DNA was transfected into mouse liver, as determined both 20 h and 7 d after HDI. Indeed, liver expression levels of several type-I interferon response genes declined significantly from 20 h to 7 d after HDI, indicating that the 7 d period from HDI until liver collection for STARR-seq reporter activity assay may allow for decay of any initial, low-level interferon response, including plasmid DNA-independent interferon responses associated with HDI itself. The 7 d period was also sufficient (Fig. S3) to resolve the initial liver histopathological responses seen in the hours following HDI [75].

### Scalability of HDI-STARR-seq and its limitations

HDI-STARR-seq is flexible and can be scaled up by cloning, in a one-pot reporter library synthesis, a complex pool of genomic DNA fragments released from open chromatin regions by mild DNase-I digestion of liver nuclei, similar to ATAC-STARR-seq [78, 88]. HDI-STARR-seq libraries may also be prepared using ChIP’d DNA for histone marks or TFs [89] or synthetic DNA fragments tiled across specific genomic regions of interest [88, 90], which facilitates fine mapping of active enhancer regions and enables discovery of functional motifs driving activity at individual enhancers. Increasing HDI-STARR-seq library complexity 500-fold, from that of a focused library comprised of 100 cloned inserts to > 50,000 DHS regions, gave good results, but resulted in some increase in variability of reporter recovery between livers.

Moreover, we observed quantitative differences in condition-specific enhancer activities between assay methods and across libraries of different complexities. For example, the *Cyp2b10* enhancer at DHS #50358 was consistently activated by 24 h TCPOBOP exposure across studies, but the fold increase in reporter activity ranged widely, from 245-fold (single enhancer HDI luciferase assay) to 14.9-fold (focused HDI-STARR-seq library) to 1.7-fold (global HDI-STARR-seq library). The lower calculated TCPOBOP responses seen with the HDI-STARR-seq libraries could be due to a decrease in assay sensitivity / increased signal to noise with increased library complexity. However, the demonstrated ability to perform STARR-seq with plasmid libraries containing millions of DNA inserts [78, 88] suggests that HDI-STARR-seq may also work well with plasmid libraries of much higher complexity. HDI can transfect up to ∼40% of hepatocytes in mouse liver in vivo [83], which may reflect the intrinsic preference of HDI for transfection of pericentral hepatocytes, noted above, and could limit the utility of HDI-STARR-seq for discovery of enhancers that control genes preferentially expressed in periportal hepatocytes [76, 91].

Other limitations, which apply to STARR-seq in general, include the need for extensive PCR amplification when preparing high complexity STARR-seq reporter RNA libraries for sequencing, which tends to decrease library complexity and introduce variability in the recovery of individual reporters [87]. This issue can be addressed by inclusion of UMIs to eliminate PCR amplification bias during preparation of STARR-seq reporter sequencing libraries [60]. There is also the variable effect that enhancer sequence has on reporter RNA stability, which can compromise the ability to distinguish strong from weak enhancers [92]. It should be noted, however, that enhancer activity comparisons across biological conditions, e.g., in male vs female mouse liver, are determined by the ratio of enhancer activity between conditions for each individual enhancer sequence and are therefore not directly impacted by this consideration.

### Promoter choice is critical for in vivo liver HDI-STARR-seq

Promoter selection is a key parameter that impacts the level of reporter transcription in MPRAs, including STARR-seq. Different core promoters have preferences for different cofactors [28, 93] and genomic environments [94] and for enhancers targeting genes of particular function [87]. Overall, more than 50% of enhancers tested show significant preferences for specific promoters [95]. We selected the mouse *Alb* promoter based on its ability to effect high level, sustained expression in mouse liver [59]. We used a minimal *Alb* promoter to ensure that basal reporter activity is low, in an effort to increase sensitivity for detection of weak enhancers and potentially discriminate between enhancers showing low vs high activity (e.g., Fig. 1B), albeit at the cost of limiting its utility for assaying repressive genomic sequences [96]. Importantly, RNA reporter activity read counts from the *Alb* promoter-based STARR-seq-transfected livers were high, which enabled us to discover 8,875 active enhancers from 50,332 reporter sequences tested. In striking contrast, liver RNA reporter read counts were 50 to 150-fold lower, precluding enhancer discovery when HDI was performed with STARR-seq libraries cloned in parallel but incorporating the SCP1 synthetic promoter or bacterial ORI promoters commonly used for STARR-seq [25], despite equivalent liver recovery of plasmid DNA for all 3 STARR-seq promoter libraries (Table S3A). Conceivably, *Alb* promoter sequences, but not SCP1 or ORI sequences, may prevent heterochromatinization and repression of the transfected STARR-seq plasmid DNA. Finally, given the sequence-encoded preferences that certain promoters may have for certain enhancers, mediated, for example, by interacting sets of TFs or cofactors [95, 97, 98], we can expect that the minimal *Alb* promoter sequence incorporated into our STARR-seq design will be suboptimal for some subsets of liver DHS, e.g., sex-biased enhancers with their primarily distal enhancers [40]. Consistent with this, active enhancers identified by HDI-STARR-seq showed strong enrichment for TSS-proximal sequences, in contrast to studies using ATAC-STARR-seq (which also interrogates open chromatin regions, as does our global HDI-STARR-seq library), where enrichment for TSS-distal enhancers was found [78, 88]. The preference of HDI-STARR-seq for TSS-proximal enhancers may result from sequence-encoded preferences of the minimal *Alb* promoter for TSS-proximal enhancer interactions. Further study, including evaluation of HDI-STARR-seq with other hepatocyte-specific minimal promoter sequences, will be needed to address this question, and to determine whether the *Alb* promoter contains activating motifs similar to those found in housekeeping gene promoters, which reduce responsiveness to distal enhancers [97].

### Chromatinization of HDI-transfected plasmid DNA in vivo and enrichments for endogenous chromatin state

A long-standing concern about using transfected reporter assays to assess enhancer activity is their reliance on episomal plasmid DNA in a non-physiological chromatin state. Indeed, MPRA activities of chromosomally-integrated DNA are better correlated with endogenous genomic annotations than the corresponding episomal sequences [99]. The transcriptional silencing of foreign DNA vectors, including plasmid vectors used for STARR-seq, has been a significant obstacle for the development of reliable reporter assays. Chromatinization of foreign plasmid DNA may yield heterochromatin that represses reporter activity over time, as was shown by the increase in repressive histone marks and the loss of activating histone marks when episomal vectors of bacterial origin were delivered to mouse liver by HDI, but not when the bacterial backbone sequences was removed (minicircle DNA vectors) [70]. We hypothesize that the inclusion in our HDI-STARR-seq plasmid library design of an *Alb* minimal promoter, with its binding sites for liver TFs, facilitates euchromatin formation and prevents the local spread of repressive heterochromatin marks initiated from plasmid bacterial backbone sequences. This proposal is consistent with our finding, noted above, that the *Alb*-promoter-based global STARR-seq plasmid library, but not STARR-seq libraries incorporating the synthetic SCP1 promoter, or the bacterial ORI promoter, yielded robust RNA reporter activity after HDI transfection in mouse liver. Moreover, the active enhancers identified by *Alb* promoter-based HDI-STARR-seq were highly enriched for genomic sequences with activating histone marks and deficient in repressive histone marks in mouse liver (Fig. 6A).

Furthermore, the set of enhancers active in male but not female liver was significantly enriched for genomic regions showing a male liver bias in chromatin accessibility, while the set of enhancers active in female but not male liver was significantly enriched for genomic regions showing a female liver bias in chromatin accessibility (Fig. 5E); thus, the sex-dependence of HDI-STARR-seq enhancer activity matches the sex-dependent pattern of endogenous chromatin accessibility seen in mouse liver. Together, these observations support the proposal that the HDI-transfected, episomal *Alb* minimal promoter-based STARR-seq reporter plasmid DNA acquires, over a 7-day period following HDI, epigenetic features similar to the corresponding chromosomal sequences in mouse liver in vivo, recapitulating the endogenous chromatin state and its regulation of enhancer activity.

Indeed, we observed in preliminary ChIP-qPCR studies that HDI-transfected episomal luciferase reporter plasmid incorporating the TCPOBOP-responsive active enhancer at DHS # 50358 (Fig. 1B) shows a robust TCPOBOP-stimulated increase in H3K27ac marks in mouse liver in vivo, as does the corresponding endogenous mouse liver chromatin sequence [36].

## Conclusions

HDI-STARR-seq is a flexible and scalable assay that can be used to identify condition-specific enhancers in a tissue-specific and cell-type-selective manner within an intact, complex organ in vivo. As exemplified here for mouse liver, HDI-STARR-seq combines liver-specific plasmid library delivery with the coupling of predicted enhancer sequences to a liver-specific *Alb* minimal promoter, and it supports large scale multiplexing of functional enhancer activity assays within a single mouse for enhancer identification.

Importantly, HDI-STARR-seq active enhancers are very strongly enriched for endogenous liver epigenetic signatures of active enhancers, suggesting they retain sufficient sequence information to recapitulate the essential chromatin state required for functional activity in vivo. HDI-STARR-seq can thus be expected to facilitate discovery of regulatory elements whose functional activity is critically dependent on the physiological cellular, metabolic and/or hormonal environment of an intact organ system. HDI-STARR-seq thus complements other MPRA methods developed for in vivo enhancer identification, including those using adeno-associated viral vectors for STARR-seq delivery to mouse brain [31–33]. The persistence of HDI-transfected transgene expression in liver for many weeks [59] suggests that HDI-STARR-seq will also be useful, not only for assaying global changes in enhancer activities in response to short-term drug or chemical exposure, as shown here using the CAR agonist ligand TCPOBOP, but also longer-term, during the course of liver disease development, e.g., following hepatotoxin exposure or when feeding high fat dietary regimens that induce steatohepatitis over a period of many weeks. HDI via the tail vein is effective for liver-specific plasmid delivery in a variety of species, including rat, rabbit and pig, and it has been adapted for regional injection supporting selective delivery to many other organs, including kidneys, muscle and pancreas in both small and large animals [100]. HDI-STARR-seq can thus be adapted to characterize enhancer activities in a variety of species and tissues by selecting suitable tissue-specific and species-specific promoter sequences.

## Ethics approval

All mouse work was carried out in compliance with procedures specifically reviewed for ethics and approved by the Boston University Institutional Animal Care and Use Committee, protocol PROTO201800698, and in compliance with ARRIVE Essential guidelines 2.0 [55].

## Availability of data

Raw and processed data for next generation sequencing datasets generated in this study are available from GEO at (https://www.ncbi.nlm.nih.gov/geo/) under accession numbers GSE267041, GSE267046 and GSE267075 (SuperSeries GSE267205). Derived datasets are provided in the Supplemental tables available on-line. Custom R scripts use for data analysis are available from the authors upon request.

## Competing interests

The authors declare that they have no competing interests.

## Funding

This work was supported in part by NIH grants DK121998 and ES024421 (to DJW).

## List of abbreviations

CAR: constitutive androstane receptor, gene Nr1i3
ChIP-seq: chromatin immunoprecipitation
DHS: DNase-I hypersensitive site(s)
ES: enrichment score
H3K27ac, H3K4me1, and H3K4me3: activating histone H3 acetylation (ac) and methylation (me) marks
H3K27me3 and H3K9me3: repressive histone H3 trimethylation marks
H3K36me3: transcribed region histone H3 trimethylation mark
HDI: hydrodynamic tail vein injection
MACS2: Model-based Analysis of ChIP-Seq analysis, v2
MPRA: massively parallel reporter assay
STARR-seq: self-transcribed active regulatory regions sequencing; Super Core 1 promoter (SCP1)
TCPOBOP: 1,4-Bis-[2-(3,5-dichloropyridyloxy)]benzene

## Authors’ contributions

T.Y.C .and D.J.W. jointly conceived of the study and developed the experimental design. T.Y.C. carried out all of the wet laboratory experiments, including animal work, cloning of STARR-seq library and Illumina sequencing sample preparation. T.Y.C. also processed the raw sequencing files, wrote the scripts required for data analysis and performed a majority of the primary data analysis. Additional data analysis was carried out by D.J.W. Preparation of figures and tables for publication were jointly carried out by T.Y.C. and D.J.W. T.Y.C. and D.J.W. jointly drafted the manuscript and D.J.W. edited and revised the final manuscript for publication. Both authors reviewed and approved the final manuscript.

## Acknowledgements

The authors thank Dr. Christine Goldfarb for assistance with mouse injections and Dr. Maxim Pytakov for assistance with scripts for enrichment analyses.

## Supplementary figures

**Fig. S1.**
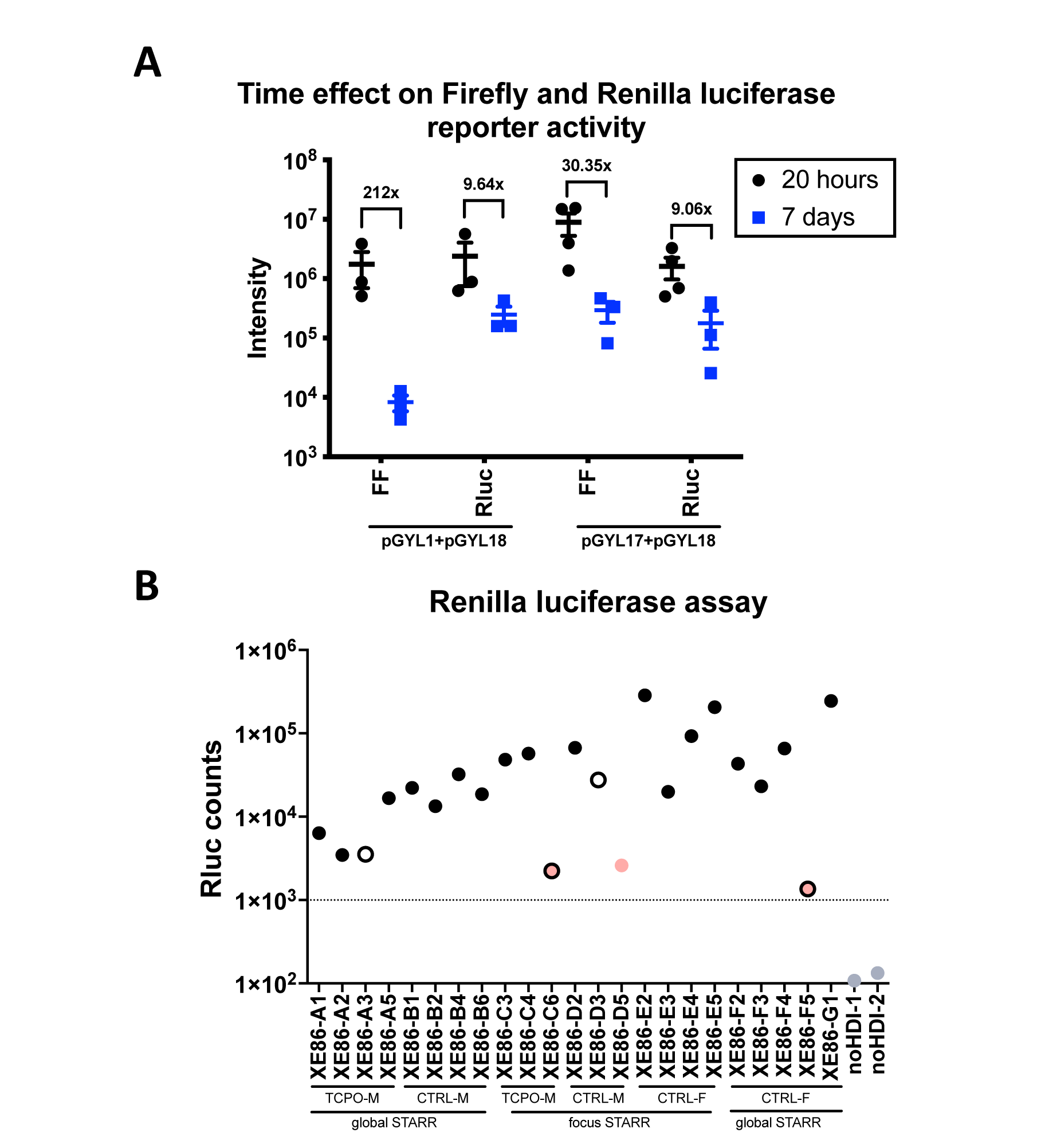
Firefly and Renilla luciferase activity in liver extracts following hydrodynamic injection (HDI). **(A)** Mice were injected with 3 μg of the internal control plasmid encoding Renilla luciferase and 15 μg of reporter plasmid (pGYL1 or pGYL17, see Table S1E) encoding Firefly luciferase by HDI. Livers were collected and assayed for firefly (FF) and Renilla (Rluc) luciferase activity at two time points: 20 h and 7 d after HDI. Data represent mean +/- SEM. Shown are the changes in each activity over time for mice injected with either a mixture of pGYL1 + pGYL18 plasmid DNA, or pGYL17+ pGYL18 plasmid DNA. Luciferase activity consistently decreased from 20 h to 7 d post-injection. The firefly luciferase activity from pGYL1 showed a greater decrease from 20 h to 7 d (212-fold) than that of pGYL18 (30-fold), while Renilla luciferase activity showed a similar decrease (∼9- fold). Each data point represents one mouse liver. See detailed data in Table S2B. **(B)** Of 36 mice, twenty-three mice were successfully injected with ∼3 ml of injection solution. 4 mice (marked as hollow circles) experienced slight resistance at the very end of the 3 ml injection, which often resulted in lower luciferase activity, as shown. Based on these luciferase activity data, we excluded from further analysis three of the lowest liver samples (marked as pink dots; XE86-C6, XE86-D5 and XE86-F5), whose Renilla luciferase activity ranged from 1,358 to 2,605; these could represent mice with poor HDI injections. Background signal, based on liver extract luciferase activity from mice without HDI transfection is marked by gray circles (background activity ∼100).

**Fig. S2.**
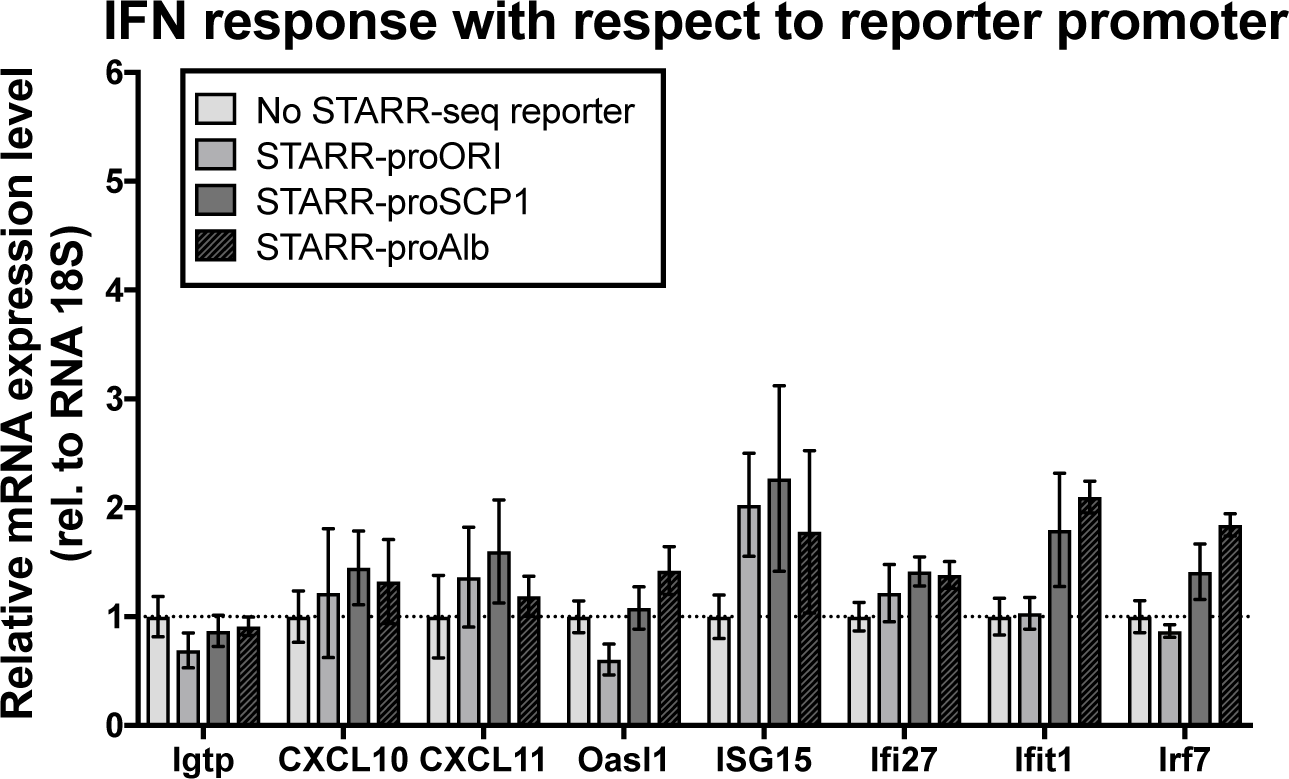
qPCR assay on Interferon (IFN)-related genes in livers of mice injected with STARR-seq reporter plasmids. Mice were injected with vehicle control (No STARR-seq reporter, plasmid pGYL18 only), STARR-seq vector with ORI promoter (STARR_proORI; STARR-TYC5 +pGYL18), STARR-seq vector using SCP1 as a promoter (STARR_proSCP1; STARR-TYC4 + pGYL18), and STARR-seq vector using *Alb* minimal promoter (STARR_proALB; STARR-TYC6 + pGYL18). See Table s1E for plasmid details. Shown are average gene expression for the indicated interferon-related genes, as determined by qPCR of extracted liver RNA. No significant difference was found in comparisons to the No STARR-seq reporter control (*t*-test, n=3-4). Livers were harvested 7 d after HDI, at which time hepatocytes recover from initial damage induced by HDI [75]. Data represent mean +/- SEM. The level of gene expression in the No STARR-seq reporter group (HDI vehicle-injected) was set = 1 for normalization. 1

**Fig. S3.**
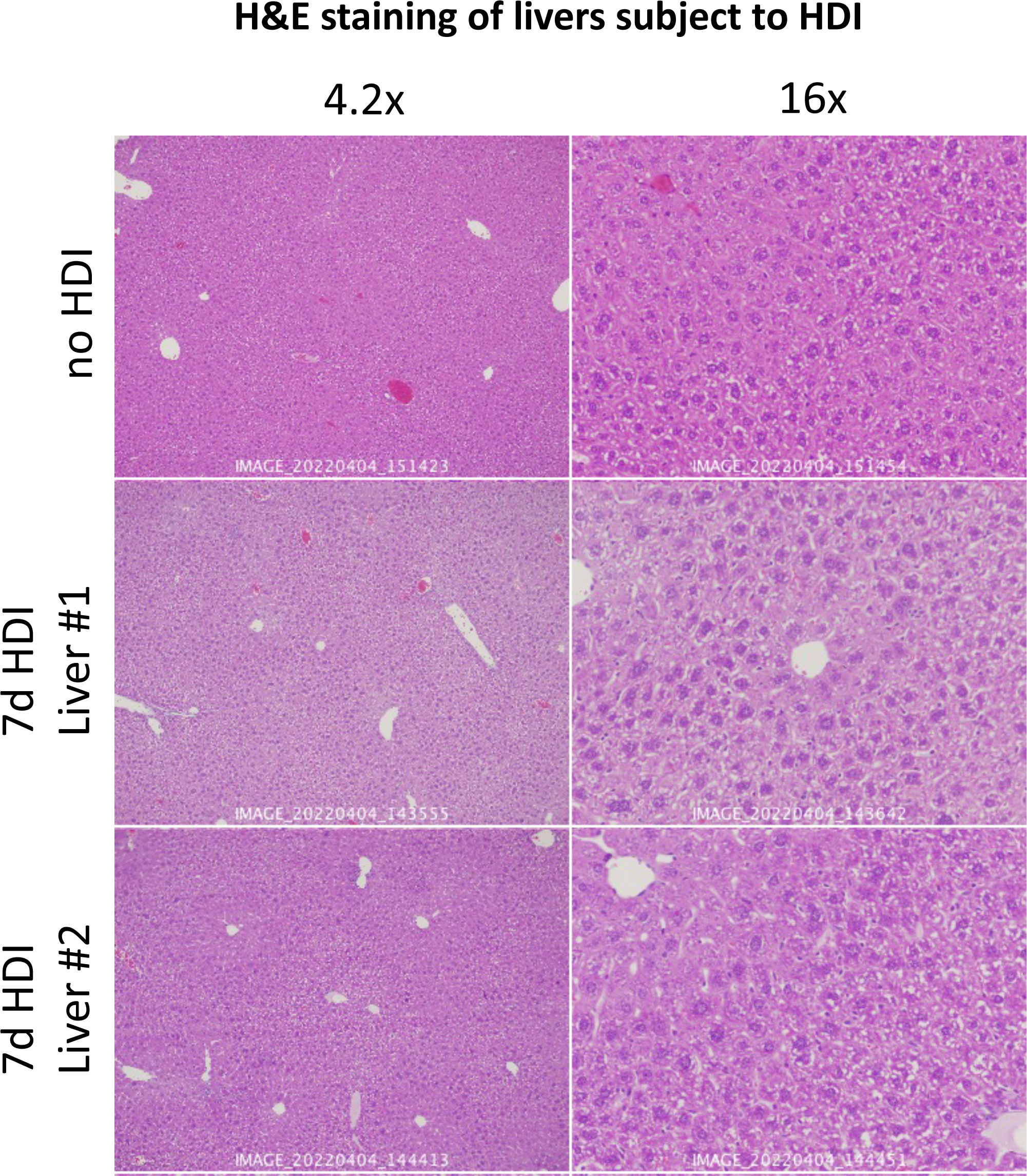
H&E staining of livers subject to HDI. Shown are liver sections from one untreated male mouse (no HDI) and two HDI-treated male mice injected with 60 μg of naked DNA plasmid (pGYL18) then stained with H&E. No histopathology was evident in mice given HDI injection after 7 days. Images were obtained using an Olympus FSX-100 instrument at 4.2× and 16×. H&E, hematoxylin and eosin.

**Fig. S4.**
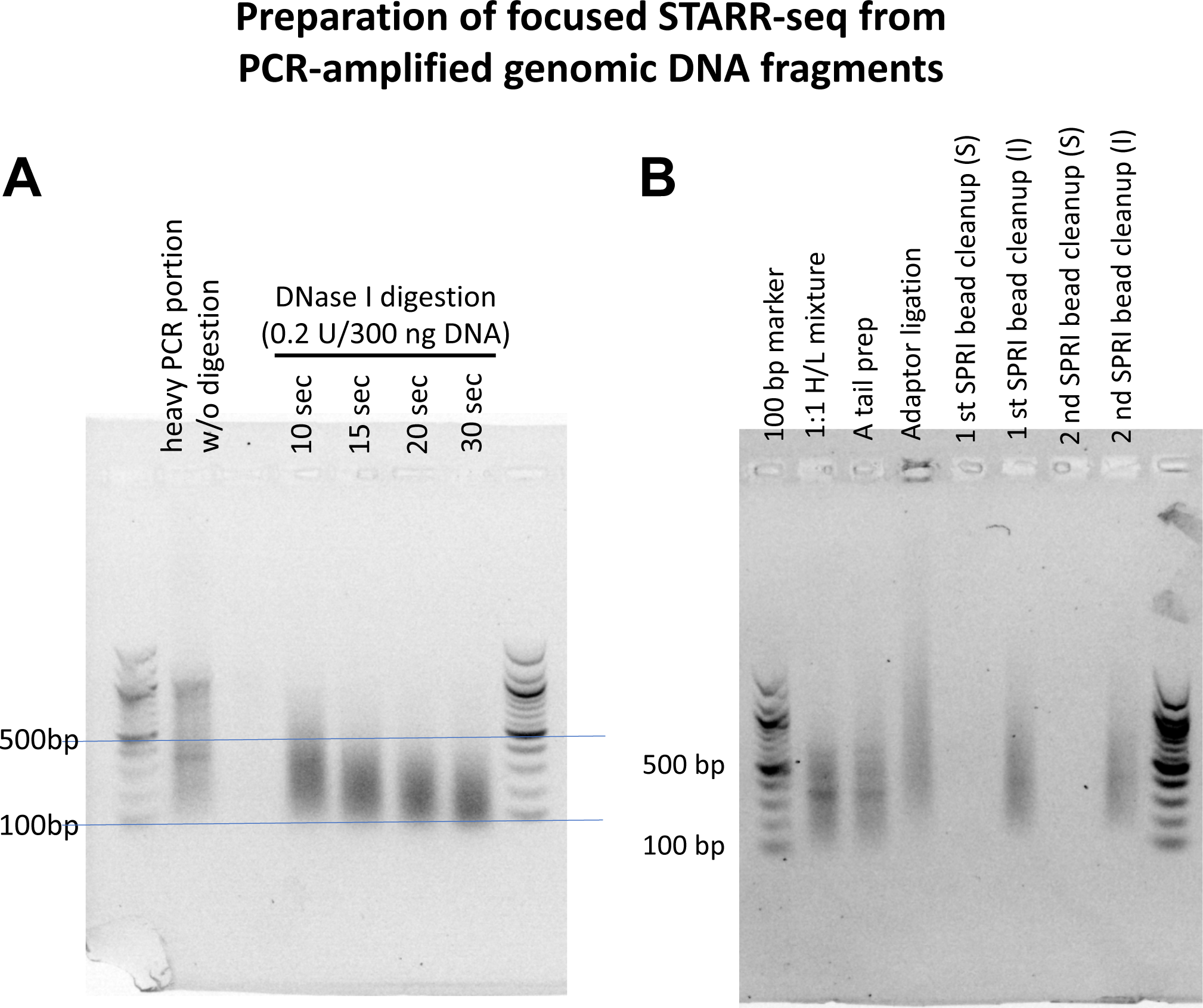
Preparation of focused STARR-seq library from PCR-amplified DNA fragments. (A) DNase-I digestion of the heavy portion of genomic DNA PCR products (sizes > 600 bp) under various conditions, as marked. We chose the 30 sec DNase-I digestion time with 0.2 U DNase-I per 300 ng DNA for the focused STARR-seq library. After DNase-I treatment, the digested larger PCR fragments ranged in size from 100 to 500 bp. (B) Size distribution of the pooled PCR-amplified DNA fragments during STARR-seq library preparation. The mixture of the smaller fragments (sizes < 600 bp) and the digested heavy PCR fragments were attached with Illumina adaptor sequences. The insert size range for the final focused STARR-seq library was 100-600 nt.

**Fig. S5.**
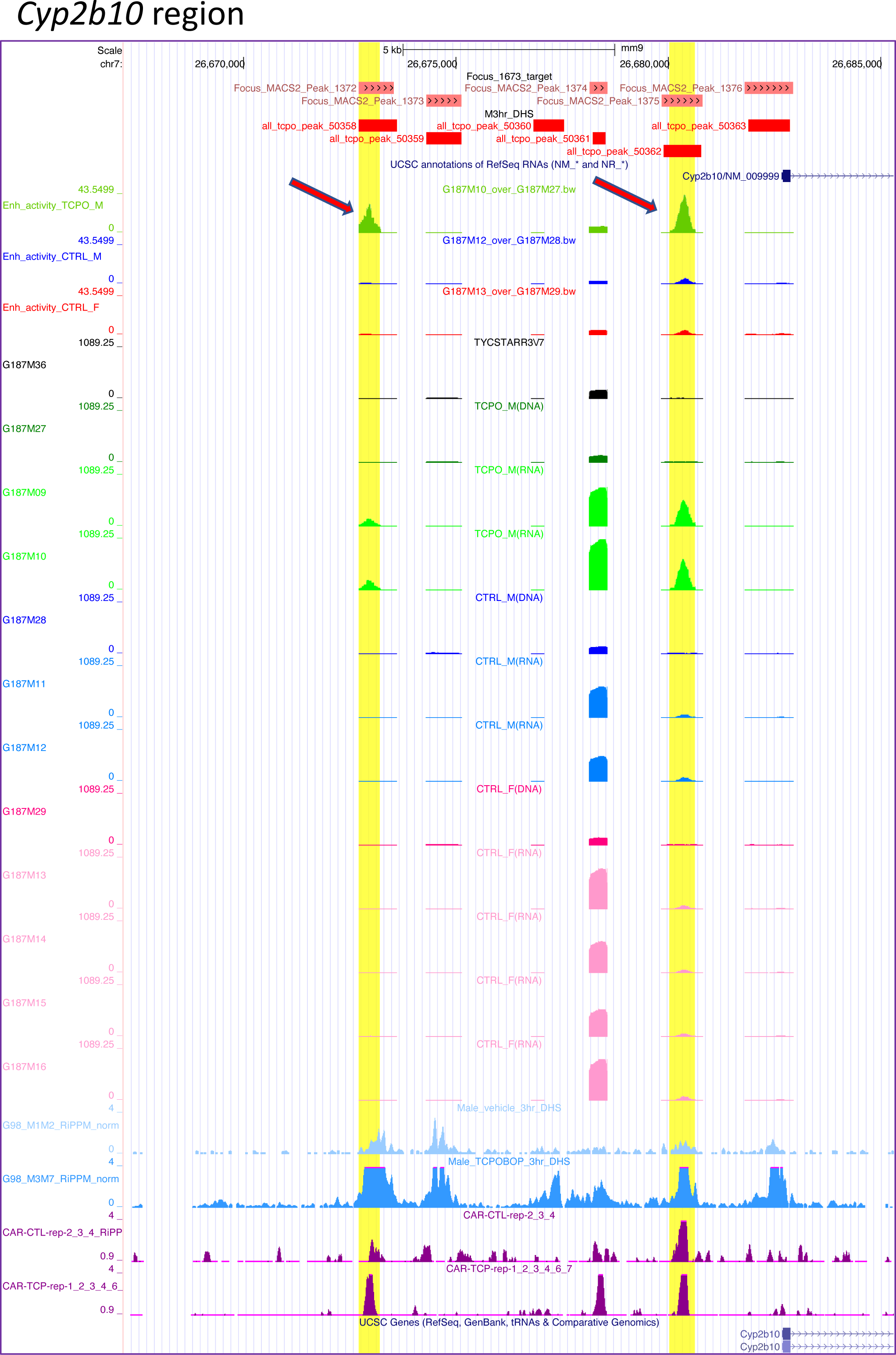
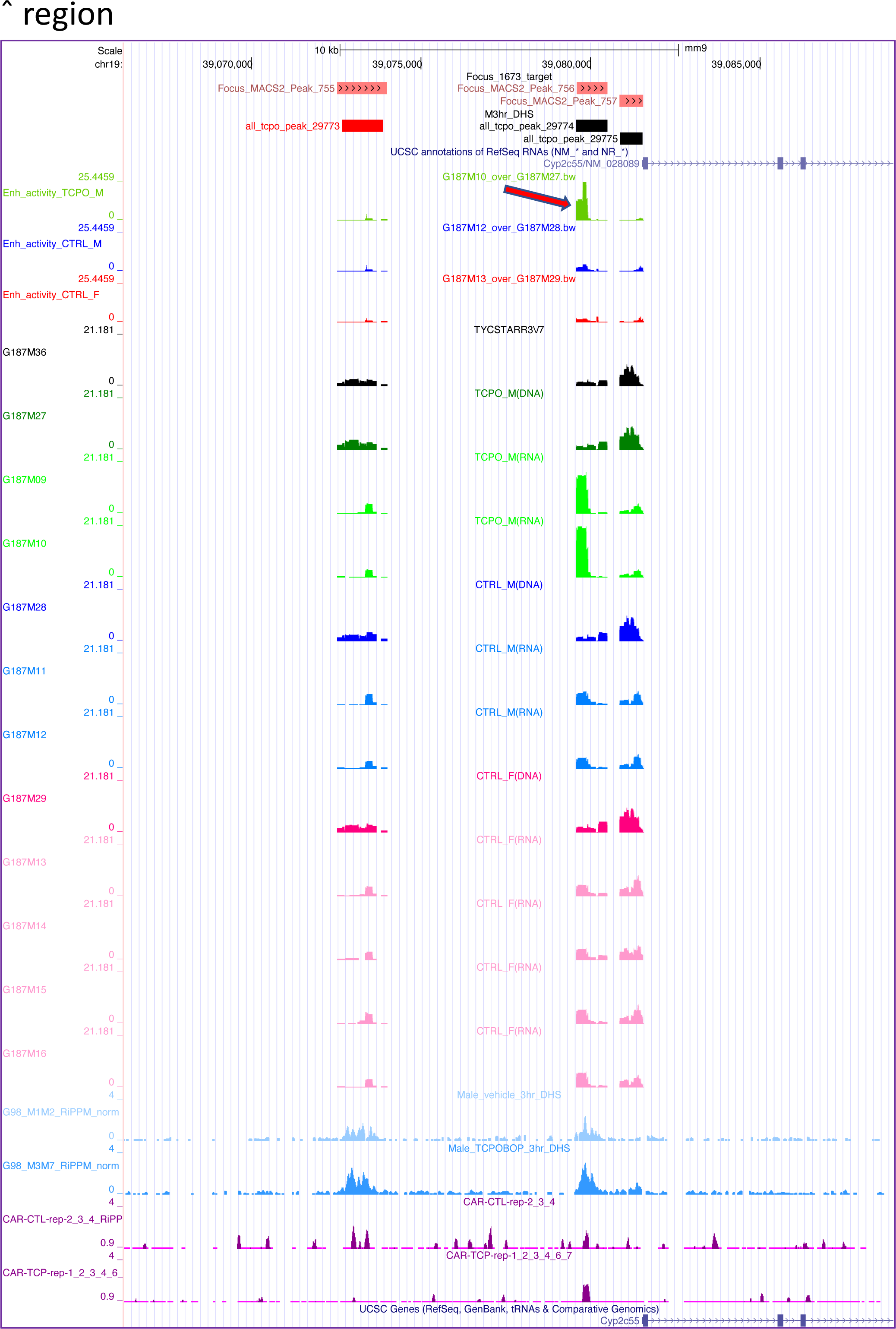
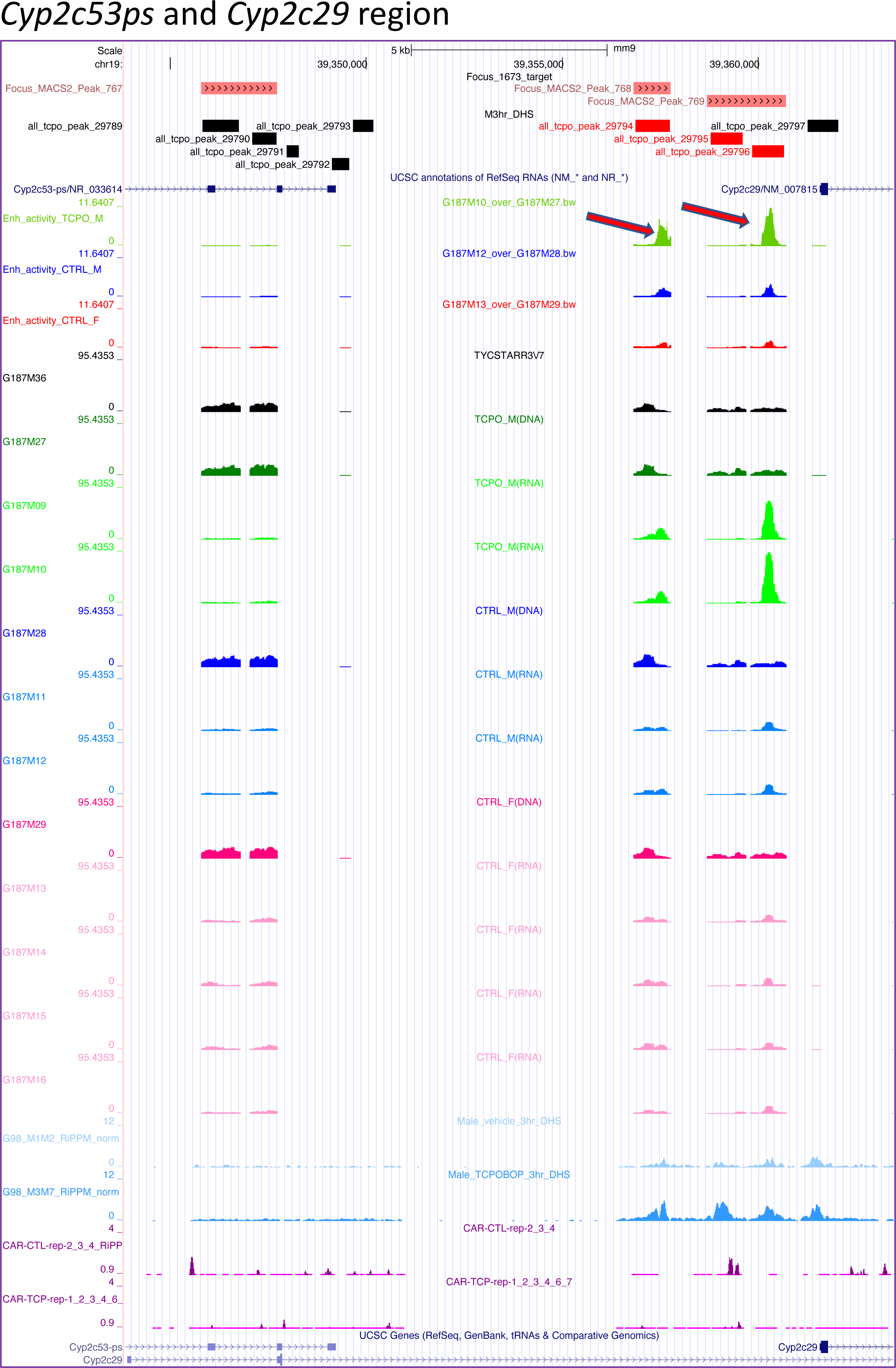
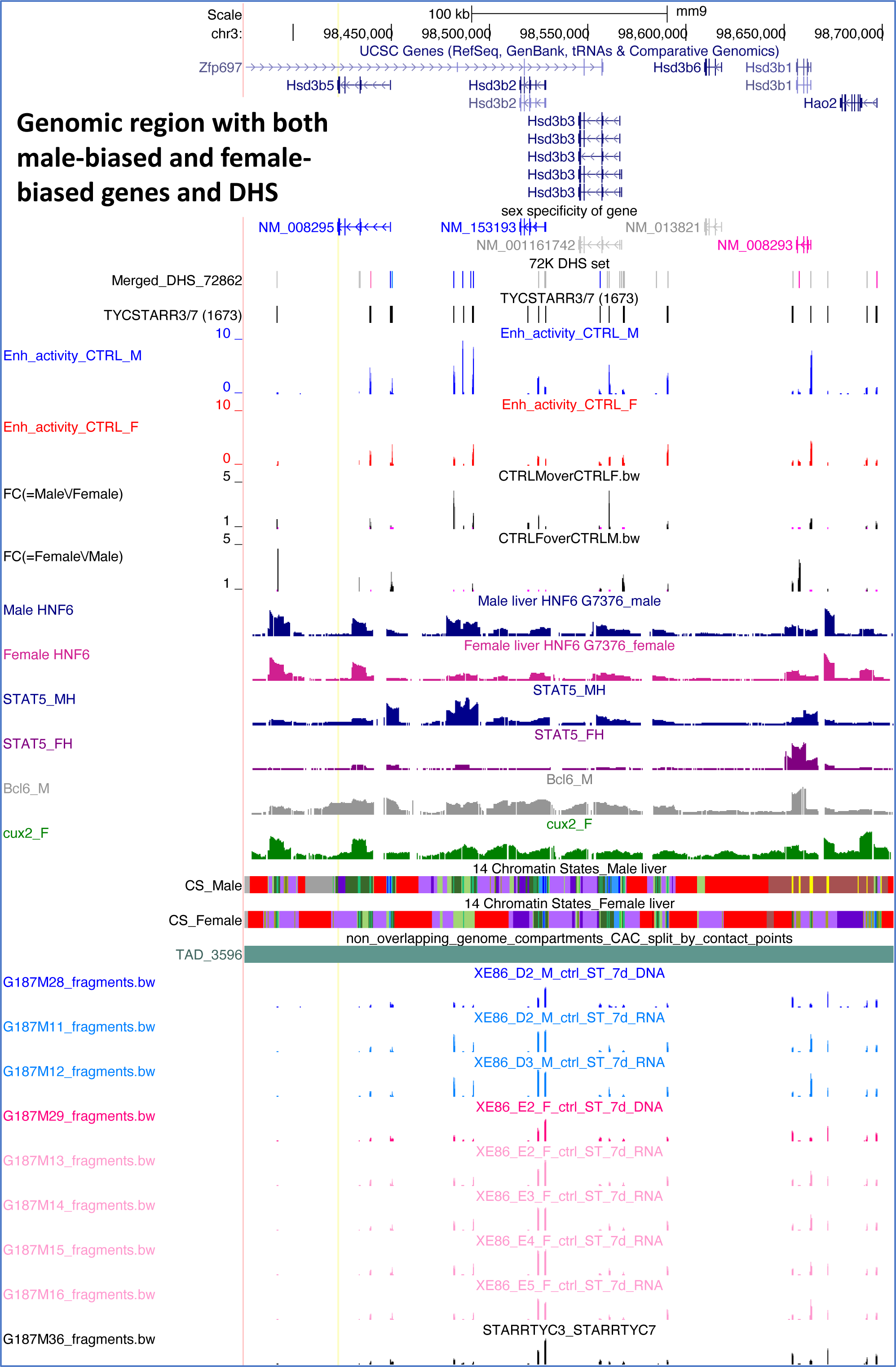
UCSC Browser screen shots: Genomic regions with TCPOBOP-responsive genes and enhancers (A-C) and *Hsd3* genomic region (D). In A-C, genomic regions showing TCPOBOP-induced HDI-STARR-seq enhancer activity are marked with red arrows. In D, the *Hsd3* region includes both male-biased and female-biased DHS and genes within the same topologically associating domain (TAD, *green bar*). UCSC Genome Browser tracks presenting STARR-seq enhancer activity at DHS sites in the vicinity of sex-biased genes *Hsd3b5* and *Hsd3b2* (male-biased expression) and *Hsd3b1* (female-biased expression). Shown are tracks for: HDI-STARR-seq enhancer activities (‘Enh’) determined in livers from male and female mice; the ratio (‘FC’, fold-change) of HDI- STARR-seq activity in males vs. females, and in females vs. males; the presence of TF binding sites, determined by ChIP-seq, for HNF6 in male and female liver, STAT5 in male and female liver, Bcl6 in male liver, and Cux2 in female liver;, and HDI-STARR-seq DNA and RNA sequence reads from 2 individual male livers (blue tracks) and from 4 individual female livers (pink tracks). Also see Table S3E.

**Fig. S6.**
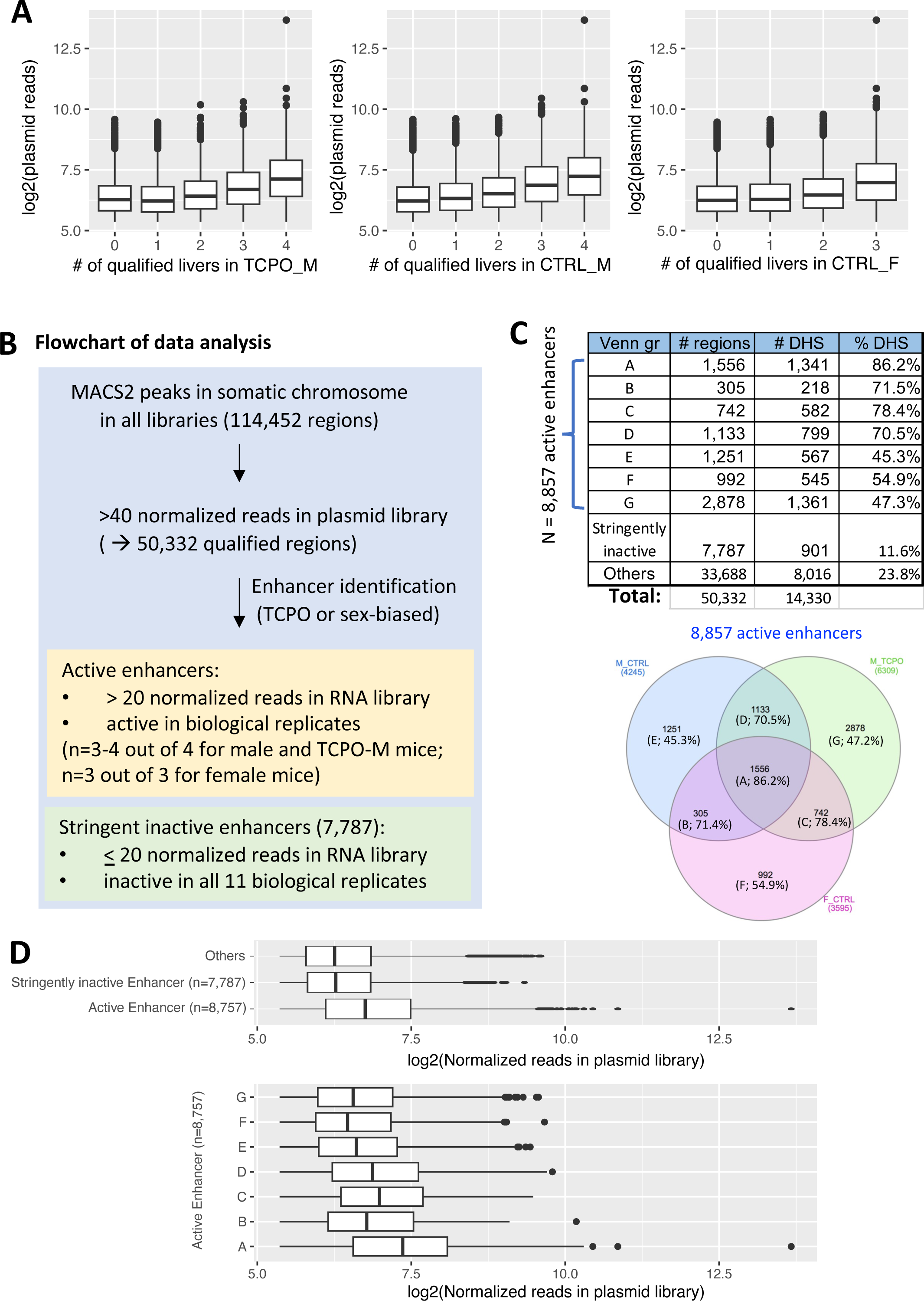
Characteristics of global STARR-seq library, prepared from DNase-I digested DNA fragments. (A) Box plots showing distributions of plasmid library reads for those RNA regions that were qualified in either 0, 1, 2, 3, or 4 of the 4 male livers analyzed (control or TCPOBOP-injected livers), or in 0, 1, 2, or 3 of the 3 female livers analyzed (bins along X-axis), where ‘qualified’ indicates the liver met the threshold of 20 normalized reporter RNA sequence reads per 10 million mapped STARR-seq RNA reads recovered from the HDI- transfected livers. Thus, STARR-seq reporter regions whose recovered liver RNA met the threshold for qualification in all 3 replicate livers (females) or in all 4 replicate livers (males) have the highest representation in the original STARR-seq plasmid library used for HDI. (B) Flowchart with thresholds used to identify active enhancers and stringently inactive enhancers. (C) Active enhancers identified by HDI-STARR-seq in mouse liver. Venn diagram (as in Fig. 5A) indicates numbers of active enhancers identified in male (control) liver, female (control) liver, and TCPOBOP-treated male liver. A total of 1,556 enhancers (Group A) are designated robust, as they show activity under all 3 biological conditions. Numbers in parenthesis presents the percentage of genomic regions in each set that overlaps a DNase hypersensitive site (also see table, above). (**D**) Boxplots showing distribution of plasmid reads in log2 scale for the set of 8,857 active enhancers (union of 4,245, 3,595 and 6,309 active enhancers identified in control male, control female and TCPOBOP-treated male, respectively), and for the set of 7,787 stringently inactive enhancers, and for each of the active enhancer sets defined in panel C.

**Fig. S7.**
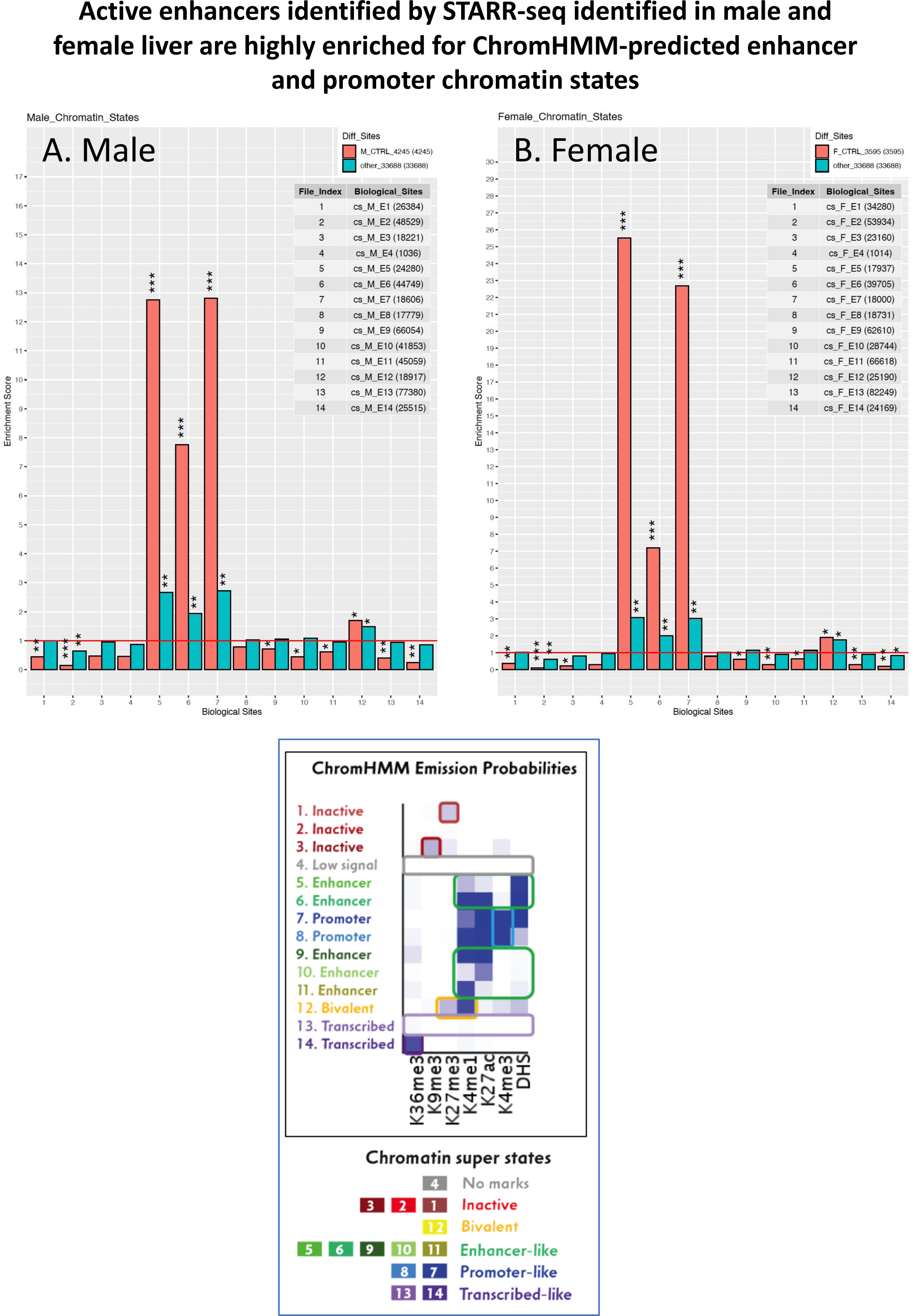
Active enhancers identified by HDI-STARR-seq are highly enriched for enhancer-like and promoter- like chromatin states predicted by ChromHMM analysis for male and female livers. Bar plots show the enrichment or depletion of each of 14 chromatin states, defined for male liver (A) or defined for female liver (B), in the sets of 4,245 HDI-STARR-seq enhancers that show activity in male liver (A) or the 3,595 enhancers that show activity in female liver (B). Enrichment scores (Y-axis) are shown for the indicated sets of STARR-seq active enhancers (blue-green bars) and for the full set of 50,322 qualified regions included in the global STARR- seq library (red bars). Enrichment scores were computed relative to a background set of 7,787 stringently inactive genomic regions. The significance of enrichment was determined by Fisher’s exact test: *, P<0.001; **, P<1e-10; ***, P<1e-100. *Bottom*: Emission probabilities for each of the 6 histone marks and DHS used to define the 14 chromatin states predicted by ChromHMM in male liver, and separately, in female liver.

**Fig. S8.**
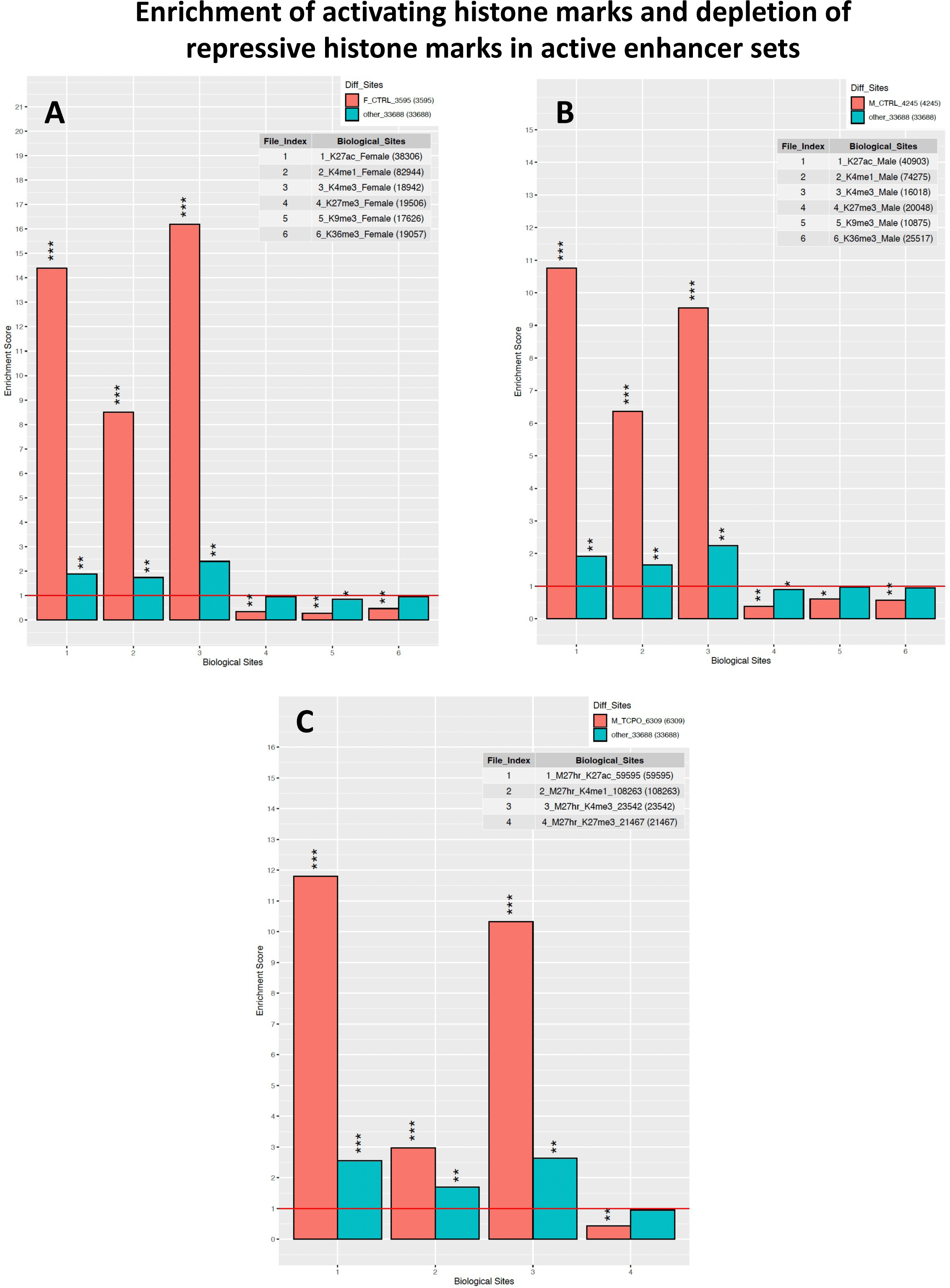
Active enhancers identified by HDI-STARR-seq are highly enriched for activating histone marks and depleted of repressive histone marks. Bar plots show enrichment scores of the indicated sets of histone marks at HDI-STARR-seq active enhancers identified in female liver (n=3,595) (A), male liver (n=4,245) (B) and in 7-day TCPOBOP treated male liver (n=6,309) (C). Comparisons were made to histone marks identified by ChIP-seq analysis of female (A), male (B) and 27 h TCPOBOP-treated male livers (C). Enrichment scores (Y-axis) are shown for the indicated sets of HDI-STARR-seq active enhancers (blue-green bars) and for the full set of 50,322 qualified regions included in the global STARR-seq library (red bars). Enrichment scores were computed relative to a background set of 7,787 stringently inactive genomic regions. The significance of enrichment was determined by Fisher’s exact test: *, P<0.001; **, P<1e-10; ***, P<1e-100.

**Fig. S9.**
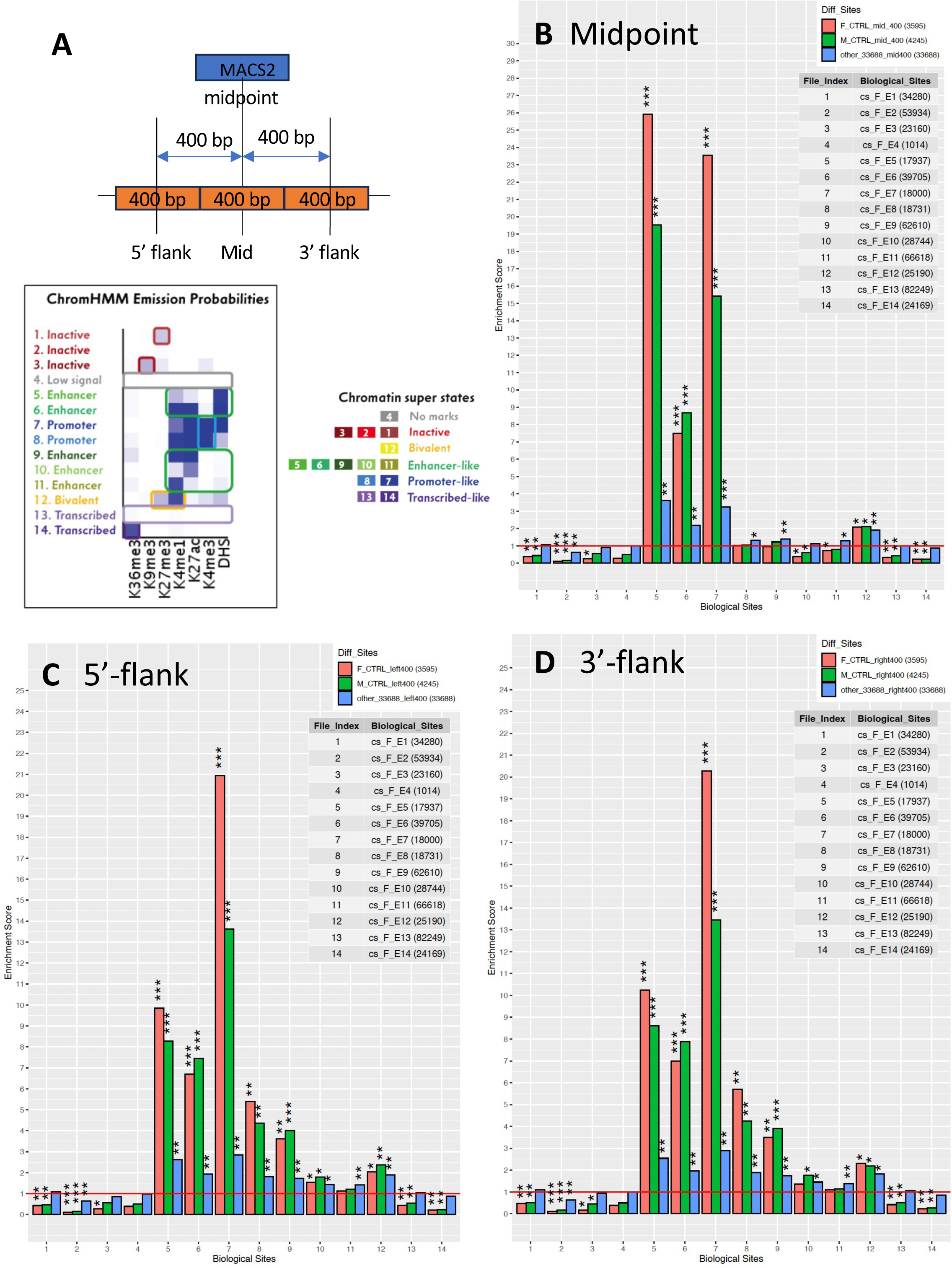
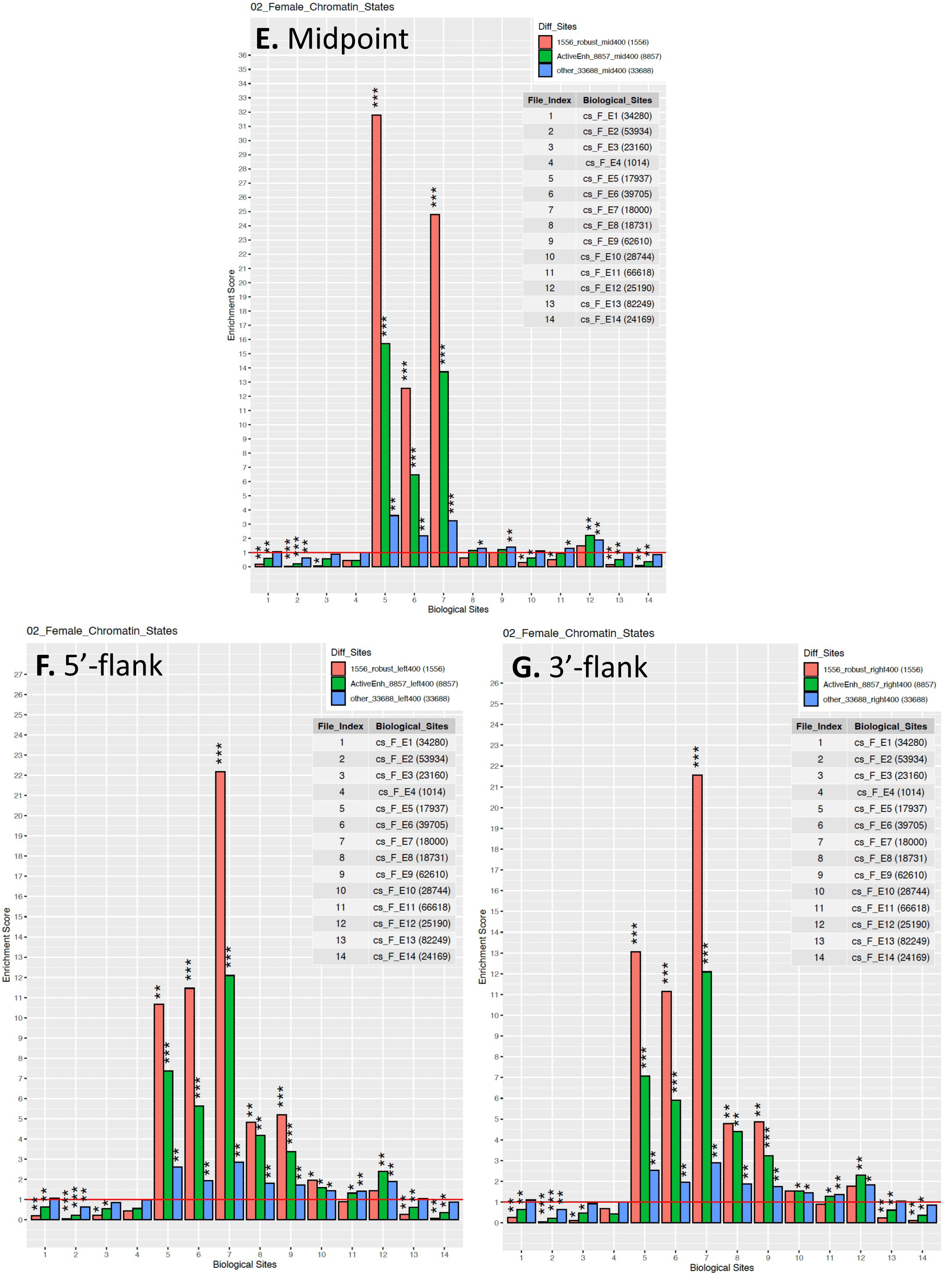
Central regions of the active HDI-STARR-seq enhancers show high overlap with enhancer and promoter features. (A) Top, schematic illustration depicting the delineation of *left*, *center*, and *right* sub- regions (each 400 nt long) relative to the midpoint of each STARR-seq active enhancer region. Below, chromatin states predicted by ChromHMM in female liver. (B, E) Enrichment scores of the central subregions surrounding the midpoint of the STARR-seq active enhancers, indicating strong enrichment of chromatin states E5, E6 and E7 (enhancers and promoters). (C, D and F, G) Enrichment scores of the 5’-flank and 3’-flank subregions, respectively, relative to the midpoint of the active STARR enhancers, indicating strongest enrichment for chromatin state E7, a promoter state. Data in panels B-D are for the sets of active enhancers identified in female (red) and male liver (green); data in panels E-G are for the sets of 1,556 robust active enhancers (red) and the full set of 8,857 active enhancers active under at least one biological condition (green). Blue bars: data for the set of 33,688 other sequences in the library that show enhancer activity in at least one of 11 of the individual livers tested but are not sufficiently reproducible between livers to meet the threshold for active enhancers. Enrichment scores were computed relative to a background set of 7,787 stringently inactive genomic regions. The significance of enrichment was determined by Fisher’s exact test: *, P<0.001; **, P<1e-10; ***, P<1e-100.

**Fig. S10.**
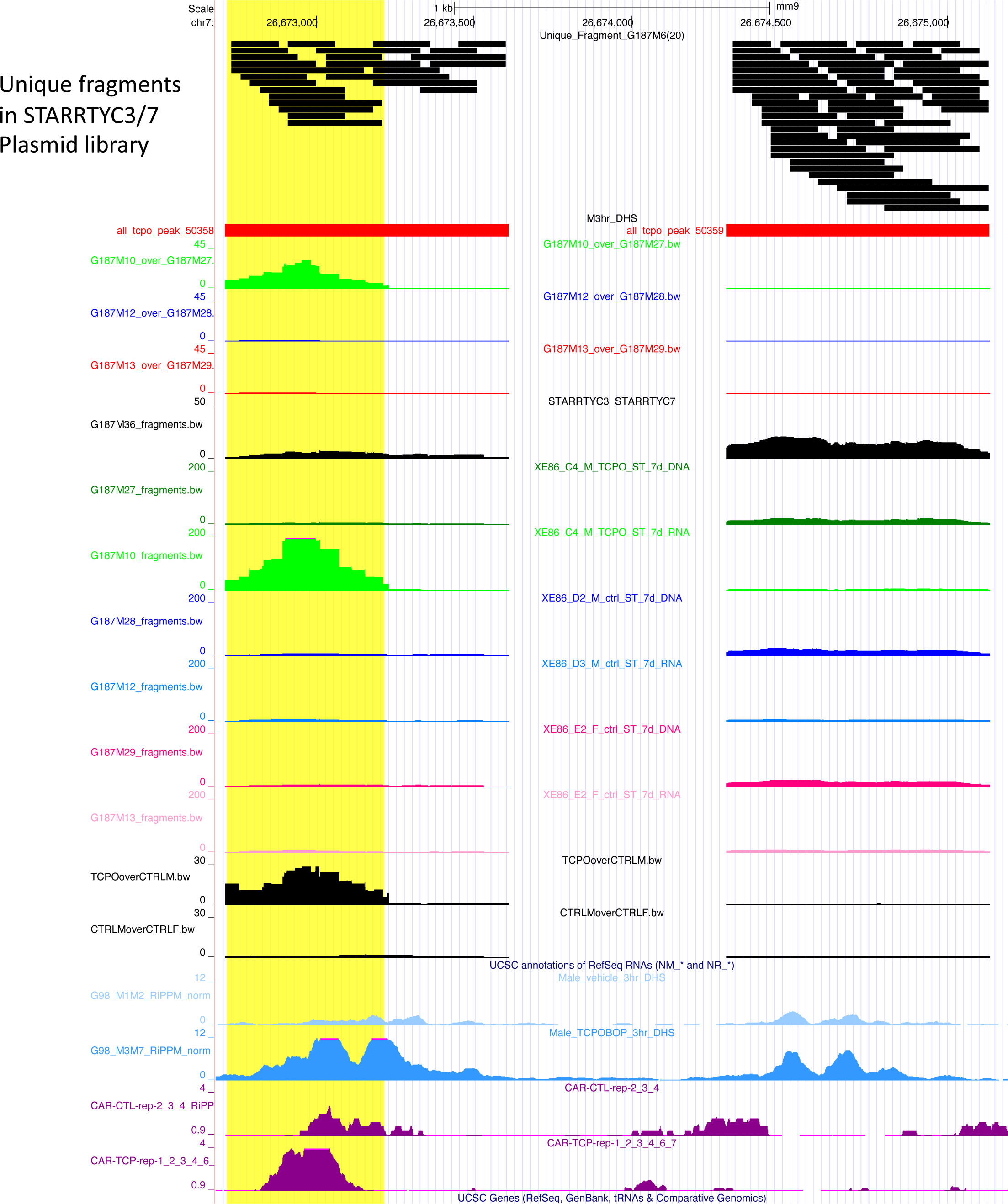
Example of unique fragments in focused STARR-seq library TYC3/7. Black horizontal bars in track at the top provide examples of 2 of the 100 cloned genomic regions represented in the focused library and show that they contain a large number of unique, cloned DNA fragments, owing to the light DNase-I digestion applied to the pooled PCR fragments mixture prior to cloning into the STARR-seq library. This diversity of reporter fragments enables refined mapping of the active enhancer sub-region within each of the 100 genomic regions, as illustrated by the normalized RNA reporter activity within the DHS sub-region highlighted in *yellow*. A total of 2,083 unique sequence fragments (unique reporters) mapped to the 100 cloned DHS regions (Table S3B, column AY). To determine the number of unique reporter sequences in each region, we first counted the number of unique reporters within each MACS2 peak. Next, to account for sequencing errors of a few base pairs, we rounded up the start and end coordinates by 10. After this rounding process, mapped fragments with unique pairs of start and end coordinates (Table S3B, column AY) were considered as unique reporters.

## Notes

### Competing Interest Statement

The authors have declared no competing interest.

